# Lineage Memory and Convergent Adaptation Dictate the Single-Cell and Spatial Architecture of Brain Malignancies

**DOI:** 10.64898/2026.06.15.732237

**Authors:** Fei Wang, Xiaodong Pang, Chen Yang, Run Huang, Wenqian Cao, Yuhan Bai, Haohao Qiu, Juyi Zhang, Bixi Gao, Chao Ma, Yanbo Yang, Shengkai Yang, Dengfeng Lu, Zhouqing Chen, Xiaoou Sun, Zhong Wang

**Author notes:** Correspondence: **Zhong Wang**, **Dengfeng Lu**, **Zhouqing Chen**, **Xiaoou Sun**. Fei Wang, Xiaodong Pang and Chen Yang contributed equally to this work.

## Abstract

How diverse cancer lineages navigate the draconian central nervous system environment—whether constrained by ancestral ontogeny or driven by convergent adaptation—remains a fundamental biological paradox. To decode this, we integrated extensive in-house and public resources to construct the largest single-cell atlas of primary and secondary brain malignancies (>550,000 cells). We demonstrate that malignant cells balance strict lineage imprinting with shared brain-adaptive programs, powerfully converging upon a dominant pan-cancer mesenchymal (MES)-like state alongside a SYT1+ pioneer subpopulation exploiting neuronal mimicry. Concurrently, the microenvironment undergoes lineage-constrained divergence. The myeloid landscape shifts from a resident-dominated architecture in gliomas to extensive blood-borne infiltration in metastases. Lymphoid responses parallel this, crowning laryngocarcinoma as an ultra-hot subtype and revealing that massive T cell influx in highly infiltrated niches is paradoxically driven into terminal exhaustion, contrasting with severe immune exclusion in cold tumors. Furthermore, stromal-vascular adaptation couples pro-angiogenic tip-endothelial activation with a dynamic opposition between structural extracellular matrix (ECM) deposition and focal proteolytic degradation. Anchored by the pan-cancer MES adaptation, we leveraged subcellular spatial transcriptomics to redefine the microenvironment around four MES-like architectural niches, mapping an evolutionary trajectory from perivascular entry to immunosuppressive stromal remodeling. Finally, projecting these architectures onto independent clinical cohort establishes the MES-S2 proliferative niche as the primary driver of severe clinical deterioration across both transcriptomic and proteomic dimensions.

## Introduction

Evolutionarily, the brain is architected as a sequestered “ecological island,” where the unique blood-brain barrier (BBB), stringent metabolic constraints, and a highly integrated neuro-vascular network constitute a high-pressure environmental selection system^1,2^. Clinically, this niche is challenged by both intrinsic neuroectodermal malignancies and extrinsic metastatic invaders that colonize across systemic boundaries^3^. Recent genomic evidence indicates that brain metastases (BrM) are not merely terminal appendages of systemic disease but distinct biological entities with independent evolutionary trajectories^4^. By accumulating private mutations and exhibiting pronounced chromosomal instability (CIN), BrMs achieve rapid phenotypic adaptation to the brain’s austere microenvironment^5^. Although these two distinct classes of malignancies arise from vastly different developmental ontogenies, they must navigate a shared set of survival imperatives upon brain colonization: evading specialized CNS immune surveillance, hijacking neuro-metabolic coupling, and remodeling the host stroma to permit parenchymal invasion^6^. Crucially, emerging evidence hints that the immune-polarization landscape of the brain tumor microenvironment may be dominantly shaped by the tumor’s lineage of origin rather than its anatomical location^7^. This “hetero-organic, co-locational” competitive landscape frames a fundamental evolutionary paradox: does hard-wired cell lineage memory (ontogeny) dictate the final malignant phenotype, or does the draconian selection pressure of the brain ecosystem (environment) compel disparate malignancies to undergo functional convergent evolution?

The phenotypic evolution of malignant cells upon brain colonization represents a fundamental negotiation between ancestral ontogeny and local environment. A critical knowledge gap remains: do heterogeneous metastatic lineages co-opt the canonical state-plasticity toolkits—such as the mesenchymal (MES) or neural-like transitions—defined in glioblastoma, and are these states organized into similar layered spatial architectures^8–10^? A more provocative hypothesis is whether “neuronal mimicry”—the hijacking of neuronal programs and synaptic genes to integrate into host circuits—is a universal “brain-adaptive” strategy across diverse cancer types^11^. Defining the boundaries between biological ancestry and environmental instruction has become the essential path to understanding malignant adaptation^12^. However, whether structurally disparate primary tumors converge on identical plastic states upon brain colonization, or follow divergent, lineage-constrained evolutionary trajectories, remains fundamentally unknown.

Microenvironmental remodeling, rather than genomic evolution alone, serves as the primary engine driving tumor trajectories^13^. In contrast to the highly immunosuppressive “immune deserts” characteristic of gliomas, BrMs often exhibit pronounced “immune-hot” phenotype^14^. This immunological heterogeneity is most striking in the myeloid compartment: BrMs are dominated by marrow-derived macrophages (MDMs) and neutrophils that drive angiogenesis and suppression, while resident microglia are frequently excluded to the tumor periphery^15^. Furthermore, BrMs infiltrate via vessel co-option, inducing endothelial and mural cells to upregulate molecules like CD276 (B7-H3), thereby erecting a dual physical and immunological barrier at the vascular niche^16,17^. However, these observations remain largely fragmented and descriptive. A unified, cross-lineage framework that systematically characterizes the microenvironmental divergence between primary gliomas and metastatic invaders—as well as the distinct TME heterogeneity inherent among different BrM origins—remains completely unestablished.

Biological functions are ultimately codified in physical space, with the tumor microenvironment organized into stable, synergistic “Cellular Neighborhoods” (CNs). Spatial omics has revealed fundamental bifurcations in the topological architecture of brain tumors: the diffuse, infiltrative patterns of gliomas versus the compact, exclusionary organization of BrMs^18^. Within the vascular niche—a core functional module—the spatial downregulation of endothelial tight-junction proteins reveals the dynamics of vessel co-option and explains the spatial correlation between barrier disruption and tissue edema^19^. Deciphering this spatial logic—the transition from disordered aggregation to ordered niches—is a prerequisite for the precision targeting of microenvironmental functional modules^1^. However, a high-resolution, multi-scale mapping that captures how specific malignant programs functionally evolve across these spatial niches—and whether such localized architectural ecosystems translate into systemic, proteogenomic clinical phenotypes—has yet to be achieved.

To address these critical gaps, the present study integrates a large-scale transcriptomic atlas of primary gliomas and pan-cancer brain metastases to delineate the fundamental logic of malignant evolution and cross-lineage microenvironmental divergence. We systematically characterize the trajectories of convergent evolution versus lineage-specific divergence across both epithelial tumor cells and their accompanying multicellular ecosystems. Furthermore, we leverage high-resolution spatial transcriptomics to resolve how specific states—most notably the MES-like program—evolve functionally in space, transitioning from perivascular colonization to deep stromal remodeling. Combined with clinical proteomic validation, our work establishes a cross-lineage co-evolutionary model of brain malignancy, providing the theoretical cornerstone for identifying precision therapeutic targets within distinct spatial niches.

## Results

### A Unified Single-Cell Atlas Resolves Distinct Lineage Imprinting between Primary and Secondary Brain Malignancies

To systematically map the multi-lineage cellular architecture of intracranial tumors, we constructed a comprehensive single-cell transcriptomic atlas comprising 554,679 high-quality cells derived from 16 primary gliomas and 157 secondary BrMs, integrating 12 independent cohorts including our in-house GusuCohort (**Figures 1A, 1G, and S1A**). This pan-cancer dataset spans primary gliomas alongside a broad clinical spectrum of BrM origins, including lung, breast, larynx, skin, ovary, colorectum, stomach, kidney, and soft tissue (**Figure 1B**). Following batch integration, unsupervised clustering resolved six major transcriptionally distinct lineages: malignant cells (296,230 cells), myeloid cells (137,909 cells), lymphocytes (86,800 cells), oligodendrocytes (17,553 cells), stromal cells (12,477 cells), and vascular cells (3,709 cells) (**Figures 1E and S1B–S1G**).

**Figure 1.**
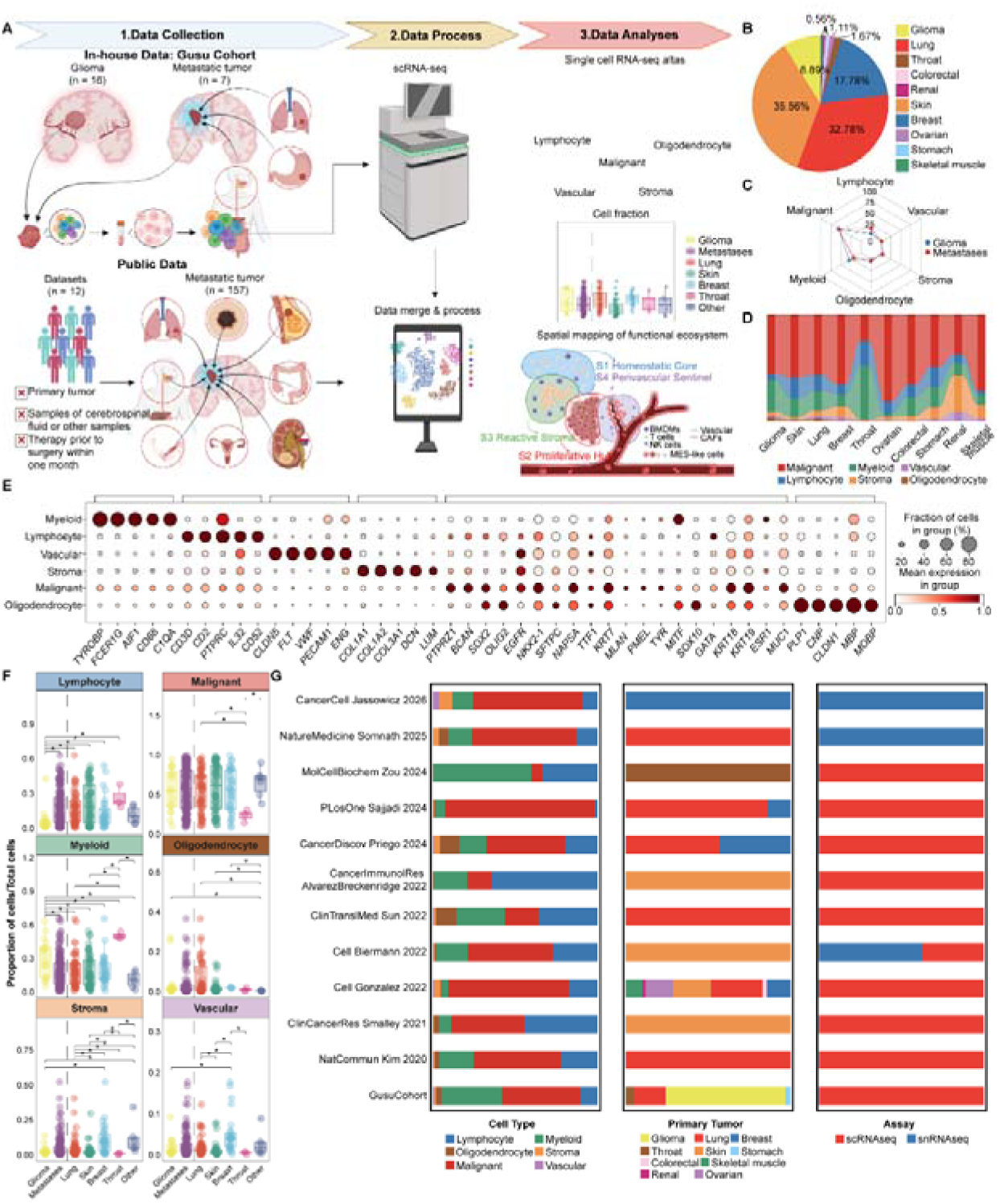
Global Multi-lineage Transcriptomic Atlas and Cellular Architecture of Primary and Secondary Brain Malignancies. **(A)** Schematic workflow illustrating study design, from clinical specimen procurement to computational batch integration. **(B)** Proportional composition of the multi-center cohort, comprising primary gliomas (n = 16) and secondary brain metastases (BrMs) stratified across nine primary tissue systems. **(C)** Sankey diagram showing individual sample cell allocation across six major lineages. **(D)** Radar chart comparing major lineage fractional abundance between gliomas and BrMs. **(E)** Dot plot showing canonical marker expression across major cell lineages. Dot size represents expressing cell percentage; color indicates row-standardized average expression. **(F)** Proportional cell type analysis across tumor cohorts stratified by primary sites. Statistical significance determined by a two-tailed Wilcoxon rank-sum test. **(G)** Dataset characterization matrix tracking single-cell composition, tumor lineages, and sequencing modalities.

Deconvolution of this global landscape unveiled a profound compositional dichotomy within the TME between primary and secondary malignancies. While malignant cells constituted the dominant population across all entities, the non-malignant immune compartments displayed highly divergent polarization patterns (**Figure 1C**). Primary gliomas exhibited a myeloid-heavy, lymphoid-depleted architecture, characteristic of an immunologically cold phenotype. Conversely, secondary BrMs demonstrated prominent lymphoid infiltration alongside a contracted resident myeloid compartment, signifying an immunologically active TME (**Figure 1C**). Stratification by primary tumor sites further unveiled distinct lineage-specific infiltration tracks (**Figure 1D**). For instance, laryngocarcinoma BrMs showed specialized myeloid enrichment, whereas renal cell carcinoma BrMs skewed toward stromal and vascular expansion. Comparative quantification across standardized clinical groups confirmed that this hot-versus-cold microenvironmental polarization is primarily dictated by the tumor’s primary site of origin rather than the central nervous system anatomy (**Figure 1F**). Together, these findings demonstrate that while primary and secondary brain tumors share a homotypic malignant dominance, their non-malignant ecosystems undergo divergent, lineage-constrained remodeling.

### Convergent Brain Adaptation and Lineage Imprinting Govern Malignant Cell Heterogeneity

To systematically characterize the intra-tumoral heterogeneity of malignant cells in BrM, we performed consensus non-negative matrix factorization (cNMF) on 282,815 high-quality single-cell transcriptomes. In contrast to traditional clustering, cNMF effectively identifies fluid co-expression modules with functional overlap. Optimization of the decomposition rank (k=16) maximized the balance between stability and reconstruction error (**Figure S2A**). After filtering seven confounding programs enriched with non-malignant signatures (e.g., myeloid c6/14/15/16 and lymphoid c11/12/13; **Figure S2B**), we identified nine biologically distinct malignant functional programs (**Figure 2A**) with robust batch effect removal (**Figures S2C–E**).

**Figure 2.**
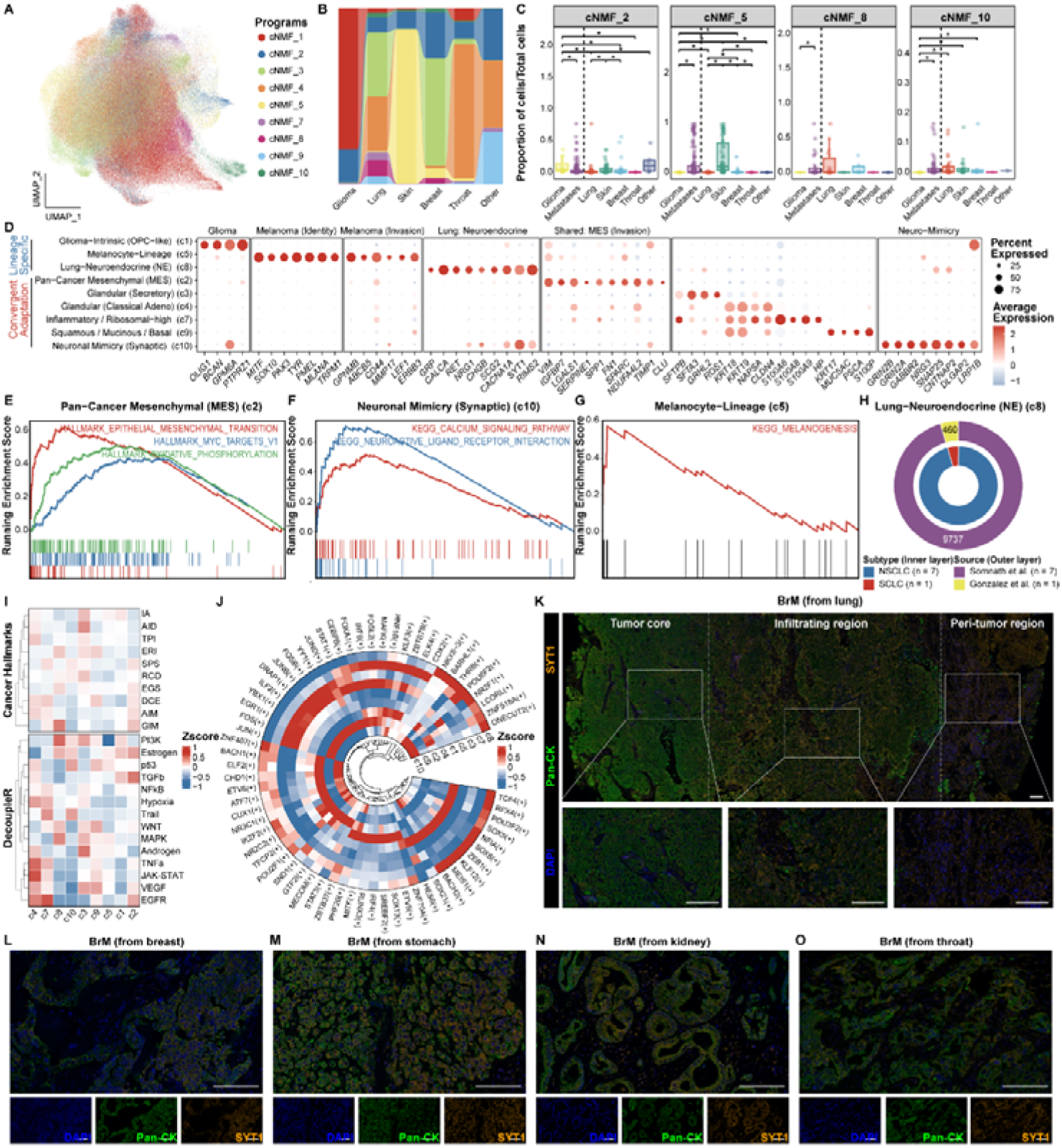
Transcriptomic Heterogeneity and Evolutionary Paradigms of Malignant Cells in Brain Malignancies. **(A)** UMAP embedding of malignant cells colored by 9 consensus Non-negative Matrix Factorization (cNMF) meta-programs. **(B)** Sankey diagram showing cNMF program distribution across primary tumor origins. **(C)** Fractional abundance box plots of key functional cNMF programs (c2, c5, c8, c10) across cohorts. **(D)** Marker gene expression profiles across cNMF meta-programs. **(E–G)** GSEA plot for the pan-cancer c2 program **(E)**, and KEGG pathway profiles for c10 **(F)** and c5 **(G)** programs. **(H)** Donut chart showing c8 program histopathology and dataset composition. **(I)** Heatmap displaying decoupleR pathway activity and cancer hallmarks for each program. **(J)** Circular heatmap of pySCENIC transcription factor regulon networks across programs. **(K)** Panoramic mIHC tile scan (DAPI, blue; PanCK, green; SYT1, orange) and regional magnification (core, front, margin) of a lung BrM section. Scale bar, 130 μm. **(L–O)** Representative mIHC validation profiles for breast **(L)**, stomach **(M)**, kidney **(N)**, and throat **(O)** metastases. Scale bar, 130 μm.

Quantification of these nine programs across glioma, lung cancer, melanoma, breast cancer, laryngocarcinoma, and other primary origins (**Figure 2B**) revealed two major evolutionary paradigms: convergent adaptation and lineage specificity. The convergent programs (c2, c3, c4, c7, c9, and c10) exhibited prevalence across multiple BrM types. Notably, c2 was the only program significantly represented across all brain malignancies, with higher enrichment in glioma than in metastases (**Figure 2C**). Programs c3 and c10 were predominantly found in lung and breast BrM, while c4, c7, and c9 were distributed across lung cancer and other metastatic origins. Conversely, lineage-specific programs (c1, c5, and c8) were restricted to distinct origins: c1 was almost exclusively glioma-derived, c5 was nearly specific to melanoma, and c8 was predominantly restricted to lung cancer BrM (**Figure 2C**). Together, these distribution patterns indicate that while BrM cells retain a “lineage memory” of their primary site, they undergo convergent evolution to acquire shared functional modules optimized for the unique “Roman” environment of the brain.

We next characterized the convergent subpopulations through marker gene integration (**Figures 2D–J**) and multi-dimensional scoring. c2 (Pan-Cancer MES] represents a pleiotropic mesenchymal hub, highly expressing *VIM*, *SERPINE1*, and *FN1*. This MES-like state is intrinsically linked to drug resistance, immune evasion, and angiogenesis, consistent with our previous glioma characterizations^20^ and independent BrM studies^5,10,12,20,21^. GSEA confirmed activation of EMT, MYC, and oxidative phosphorylation (**Figure 2E**), suggesting these cells may “educate” infiltrating myeloid-derived macrophages (MDMs) toward a pro-tumorigenic M2-like phenotype via TGF-□ and VEGFA secretion^12^, potentially enhanced by astrocyte-induced *SERPINE1*^21^. c3 (Glandular (Secretory)) and c4 (Glandular (Classical Adeno)) represent acinar-like states. c3 is distinguished by secretory features (*SFTPB*, *SFTA3*) and high MAPK/PI3K activity, whereas c4 maintains classical glandular morphology (*NAPSA*, *CLDN4*) with elevated tumor pro-inflammatory (TPI) scores and JAK-STAT activation. c7 (Inflammatory) exhibited high *S100A8/9* expression and significant granulocyte recruitment potential, facilitating early seeding of the pre-metastatic niche despite low chromosomal instability (CIN)^22^ (**Figures S2F–G**). c9 (Squamous/Mucinous/Basal) expressed *KRT17* and *MUC5AC*, with CDX2 and HNF1B activity linking it to colorectal-derived BrM. Strikingly, c10 (Neuronal Mimicry)—prominent in lung and breast BrM—highly expressed synapse-related genes (*GRIN2B*, *SNAP25*, *NRXN1*). This suggests that tumor cells not only phenotypically mimic neurons but functionally integrate into host neural circuits via glutamatergic synaptogenesis, hijacking electrical signaling to acquire survival signals and promote parenchymal infiltration^23^. This “neuromimicry” state exhibited high activation of calcium signaling and WNT hijacking (**Figure 2F**), resembling the brain-induced “MP19_Neurol-Like” state identified by Xing et al^5,23^ (**Figure S2I**).

Examination of lineage-specific programs revealed a watershed in genomic instability and pre-adaptation. c1 (Glioma-Intrinsic) (*OLIG1*, *BCAN*) exhibited significantly lower CIN scores than all BrM programs (**Figures S2F–G**), potentially explaining the lower neoantigen burden and immune checkpoint inhibitor (ICI) response rates in glioma compared to BrM^3^. c5 (Melanocyte-Lineage) (*MITF*, *SOX10*) maintained strong pigment-production programs (**Figure 2G**) even after brain colonization. c8 (Lung-Neuroendocrine) (*CHGB*, *SYT1*) served as a high-energy “engine” for lung cancer BrM, largely deriving from NSCLC samples (**Figure 2H**). This indicates that NSCLC cells acquire neuroendocrine features—typically specific to small cell lung cancer (SCLC)—upon entering the brain. Given that SCLC originates from pulmonary neuroendocrine cells^24^, this “pre-adaptation” likely facilitates interaction with host neurons without extensive remodeling. Consistent with this notion, c8 displayed the highest CIN levels (**Figures 2I/S2F–G**) and activated PI3K/MAPK pathways (**Figure 2I**), which are critical regulators of TAM-mediated colonization^25^.

Growing evidence suggests that brain metastatic cells exploit neurobiological mechanisms, such as synaptic integration and paracrine signaling, to colonize the neural parenchyma^23^, thus we explored the role of neuronal mimicry in spatial infiltration. Both c8 (Lung-neuroendocrine) and c10 (Synaptic) exhibited an “immune-cold” phenotype characterized by downregulated TNF-α and JAK-STAT activity, providing a single-cell basis for immune evasion in metabolic/synaptic subtypes. To identify a functional “pioneer” marker, we prioritized SYT1 (Synaptotagmin 1), a primary calcium sensor that facilitates fast neurotransmitter release and unconventional protein secretion—processes established as essential for driving tumor invasiveness via specific vesicle-mediated secretory pathways^26^. Consistent with this selection, *SYT1* was significantly co-expressed in both c8 and c10 programs (**Figure S2J**). mIHC validation in lung BrM sections revealed that SYT1 expression peaks at the invasive front and diminishes toward the tumor core (**Figure 2K**). While spatial zoning was less distinct in other cancer types, membrane-localized SYT1+ cells remained widespread (**Figures 2L–O**). Together, these findings indicate that SYT1 marks a “pioneer” subpopulation utilizing neuronal mimicry to drive the invasion of the normal brain parenchyma.

In summary, this transcriptomic landscape reveals that BrM heterogeneity arises from a balance of primary lineage imprinting and convergent brain adaptation. Therapeutic strategies targeting only lineage-specific antigens may induce immune escape and transition toward brain-adaptive states (c2 or c10). These data suggest that dual-targeting of lineage-specific antigens alongside brain-adaptive states—such as inhibiting MES-mediated invasion or synaptogenic communication—may be required to overcome the complexities of metastatic evolution.

### Divergent Trajectories and Functional Specialization of the Myeloid Microenvironment in Primary and Secondary Brain Tumors

To deconvolve the myeloid microenvironment across primary gliomas and secondary BrMs, we analyzed 137,909 single-cell myeloid transcriptomes, resolving five principal lineages: bone marrow-derived macrophages (BMDMs), microglia-derived macrophages (MDMs), neutrophils, dendritic cells (DCs), and mast cells (**Figures 3A and S3C**). While macrophages constituted the primary cellular infrastructure across all cohorts, DCs and neutrophils were significantly expanded in BrMs, contrasted by a relative contraction of the resident MDM compartment (**Figures 3B and 3C**). This numerical shift suggests a competitive niche displacement model driven by the progressive infiltration of blood-borne BMDMs into expanding metastatic lesions. Crucially, distinct primary lineage imprinting shapes this architecture across metastatic subtypes: lung BrMs displayed a heightened enrichment of mast cells, breast BrMs selectively retained resident microglia, and laryngocarcinoma BrMs presented a unique hyper-inflammatory profile where neutrophils emerged as the second most abundant population (**Figure 3B**). Thus, the intracranial myeloid landscape is systematically tailored by the tumor’s primary site of origin.

**Figure 3.**
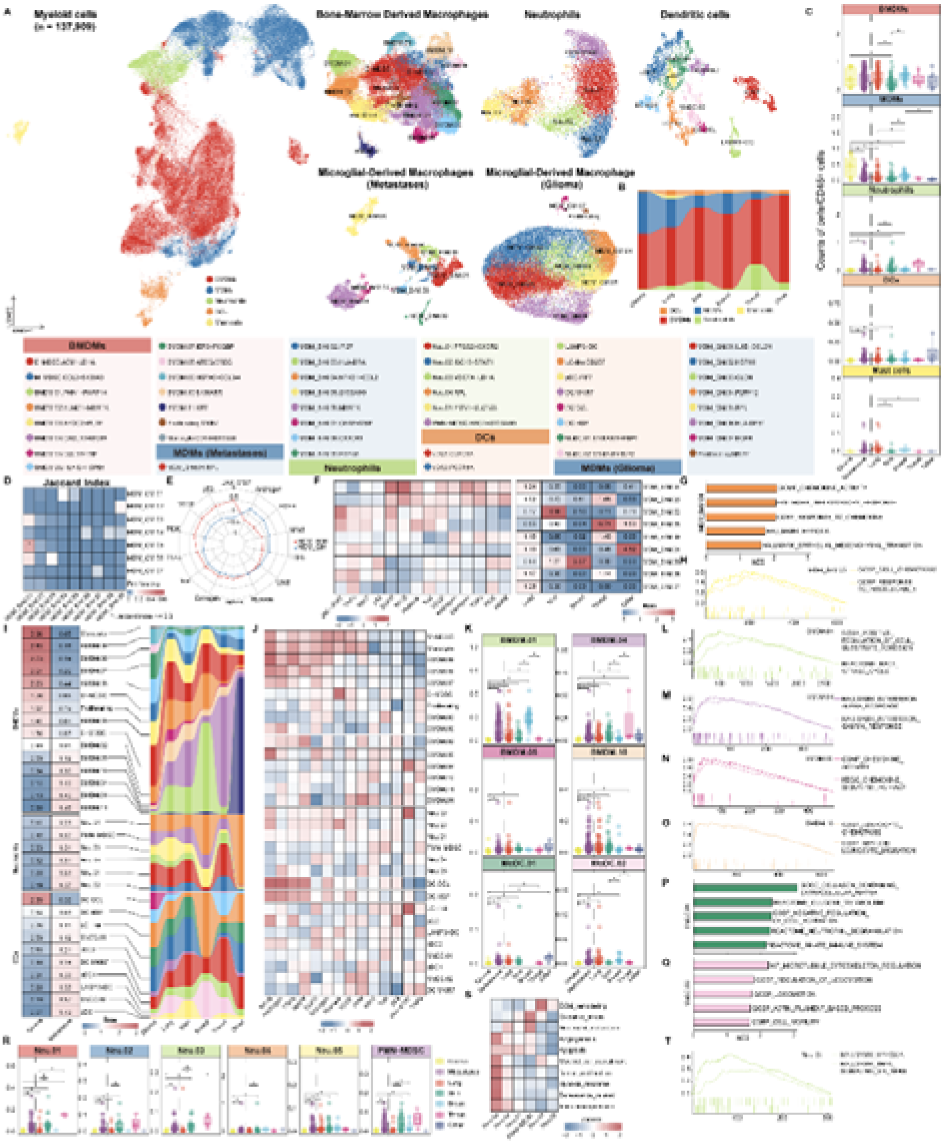
The Myeloid Landscape, Functional Polarization, and Cross-Lineage Heterogeneity in Primary and Secondary Brain Malignancies. **(A)** High-resolution UMAP of the integrated myeloid compartment. **(B)** Sankey diagram tracking myeloid lineage composition across primary tumor origins. **(C)** Fractional abundance box plots of major myeloid clusters relative to the leukocyte pool. Statistical significance determined by a two-tailed Wilcoxon rank-sum test. **(D)** Heatmap showing Jaccard similarity index across transcriptional signatures of glioma-and BrM-associated MDMs (asterisks denote index >= 0.3). **(E)** Radar chart of decoupleR signaling pathway scores in tumor-associated MDMs. **(F)** Heatmap displaying pathway enrichment (left) and observed-to-expected cell ratios (Ro/e, right) for BrM MDM sub-clusters. **(G–H)** GSEA functional terms for MG_BrM.04 **(G)** and MG_BrM.05 **(H)** sub-clusters. **(I)** Integrated analysis tracking Ro/e scores (left) and relative cohort distributions (right) for BMDM, neutrophil, and DC subsets. **(J)** Heatmap of decoupleR pathway activity split by cohort preference blocks. **(K)** Subpopulation fractional abundance comparison across cohorts. Statistical significance determined by a two-tailed Wilcoxon rank-sum test. **(L–O)** GSEA pathway curves highlighting functional phenotypes for BMDM.01 **(L)**, BMDM.04 **(M)**, BMDM.05 **(N)**, and BMDM.10 **(O)**. **(P–Q)** Functional pathway enrichment bar plots for MoDC.01 **(P)** and MoDC.02 **(Q)** subsets. **(R)** Neutrophil sub-cluster fractional abundance across cohorts. Significance evaluated by a two-tailed Wilcoxon rank-sum test. **(S)** Meta-gene activation profiles for neutrophil functional states defined by reference datasets. **(T)** GSEA tracking canonical Hallmark gene sets enriched in the Neu.03 subset.

Despite comprehensive integration, MDMs from gliomas and BrMs exhibited profound transcriptomic segregation reflecting their distinct intra-tumoral topographies; while BrM-associated MDMs are restricted to the infiltrative tumor periphery, they are ubiquitously interspersed throughout the parenchyma of primary gliomas^27,28^ (**Figures 3D and S3A–S3F**). Pathway activity profiling revealed that BrM-associated MDMs operate in a hyper-activated state, displaying elevated oncogenic signaling—including JAK-STAT, p53, VEGF, PI3K, TNF-α, TRAIL, and MAPK pathways—compared to their glioma counterparts (**Figure 3E**). These BrM-associated MDMs partitioned into three distinct functional archetypes: an angiogenic/hypoxic axis, an inflammatory/proliferative axis, and a specialized TNF-alpha-sustained module (**Figure 3F**). Phenotypically, lung BrMs demonstrated broad enrichment across all archetypes, whereas melanoma and breast BrMs selectively accumulated the MDM_BrM.06 subset, and laryngocarcinoma BrMs favored inflammatory sub-clusters (**Figures 3G and 3H**), highlighting site-specific macrophage polarization.

Characterization of the infiltrating BMDM compartment revealed a sharp programmatic bifurcation between BrM-preferred and glioma-preferred trajectories (**Figure 3I**). Glioma-preferred clusters shared a conserved network dominated by NF-kB, TGF-□, and MAPK activity, whereas BrM-preferred subsets were exclusively re-wired toward heightened JAK-STAT and TNF-α signaling (**Figure 3J**). Among the expanded BrM-preferred subsets, the gastric BrM-restricted BMDM.11 subset overexpressed epithelial-associated keratins, demonstrating tumor-coaxed phenotypic mimicry (**Figures 3I, S3G, and S3H**). Functional deconvolution showed a clear division of labor among BrM-expanded BMDM subsets: the BMDM.01 subset governed cell adhesion and trans-endothelial motility, BMDM.04 exhibited a concentrated interferon-response signature, and the BMDM.05/10 clusters regulated directed leukocyte chemotaxis (**Figures 3K–3O**). Notably, the BrM-enriched BMDM.10.S100A8/9 sub-cluster served as a potent chemoattractant hub, mirroring the pro-metastatic loops driven by S100A8+ macrophages in extracranial niches^29^.

Beyond the macrophage compartment, dendritic cells and neutrophils exhibited highly structured niche partitioning (**Figure 3I**). Glioma-preferred DCs maintained consistent activation of resident macrophage-like NF-κB and TGF-□ networks, whereas monocyte-derived DCs (MoDCs) were significantly expanded in BrMs (**Figures 3J and 3K**). Primary origins strictly dictated MoDC distribution: lung and most BrM types were dominated by MoDC.01 (specialized for ECM synthesis and glycolysis; **Figure 3P**), whereas breast BrMs uniquely enriched for MoDC.02 (dedicated to cytoskeletal remodeling and motility; **Figure 3Q**), confirming an origin-dependent functional dichotomy between migratory and antigen-presenting DC states^29^. Concurrently, neutrophil sub-clusters exhibited a near-exclusive preference for the BrM microenvironment, peaking in laryngocarcinoma BrMs (**Figure 3R**). The specialized Neu.03 sub-cluster emerged as a critical pro-tumorigenic state heavily driven by hypoxia and TNF-α/NF-κB loops, showing intense activation across modules governing immunosuppression, SASP, and microvascular angiogenesis (**Figures 3S and 3T**). These findings establish a profound compartmental dichotomy where primary gliomas enforce a suppressed, resident-dominated myeloid architecture, and secondary brain metastases undergo extensive, lineage-constrained remodeling driven by blood-borne myeloid sentinels^3^.

### Lineage-Specific Lymphoid Landscapes Reveal a Functional Dichotomy Between Cytotoxicity and Exhaustion in Brain Metastasis

To characterize the lymphoid compartment within the intracranial microenvironment, we mapped the lymphocyte lineage across diverse primary origins, resolving six major cellular identities: CD4+ T cells, CD8+ T cells, proliferating T cells, regulatory T cells (Tregs), NK cells, and B/plasma cells (**Figures 4A and 4B, Figure S4A and 4B**). T cells, particularly proliferating and Treg subsets, dominated the lymphoid pool, accounting for over 75% of lymphocytes in most niches (**Figure 4C**). Overall lymphoid infiltration was significantly elevated in secondary BrMs relative to primary gliomas (**Figure 4D**). Crucially, the immunogenic state of the metastatic niche was constrained by its primary lineage; while melanoma BrMs exhibited extensive lymphoid accumulation, breast cancer BrMs mirrored the immune-desert phenotype of primary gliomas (**Figure 4D**). Additionally, B/plasma cells were preferentially enriched in melanoma and lung BrMs, aligning with the structural presence of tertiary lymphoid structures (TLS) in these secondary lesions^30^ (**Figure 4D**).

**Figure 4.**
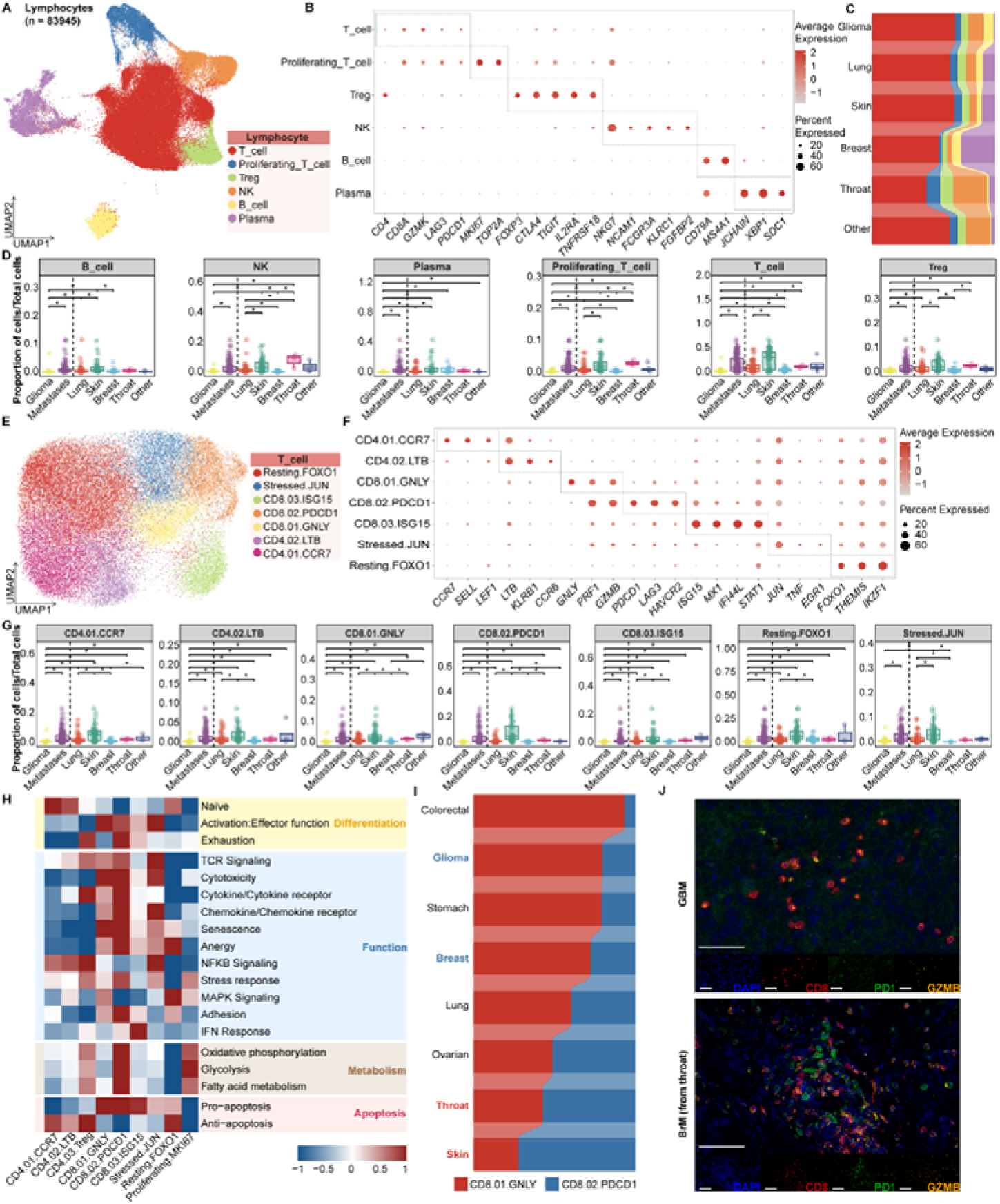
Lineage-Specific Lymphoid Landscapes and the Functional Dichotomy of T Cell States in Brain Malignancies. **(A)** Global lymphoid UMAP embedding annotated by 6 major lineages. **(B)** Canonical marker gene expression profiles defining the lymphoid lineages. **(C)** Sankey diagram illustrating lymphoid line proportions across primary origins. **(D)** Box plots of lymphoid lineage fractional abundance relative to total cells. **(E)** High-resolution UMAP sub-clustering of the non-proliferating T cell compartment. **(F)** Expression of specific marker genes across 7 T cell functional subsets. **(G)** T cell subset fractional abundance across cohorts, formatted as in **(D)**. **(H)** Heatmap grouping T cell subsets and global reference modules by functional pathway activity scores. **(I)** Sankey diagram tracking the cytolytic (CTL; CD8.01.GNLY) versus exhausted (Tex; CD8.02.PDCD1) functional dichotomy across cohorts. **(J)** mIHC profiles (DAPI, blue; CD8, red; PD-1, green; GZMB, orange) of T cell spatial infiltration in GBM (top) and laryngocarcinoma BrM (bottom). Scale bar, 130 μm.

Intriguingly, laryngocarcinoma BrMs occupied the absolute upper extremity of this immunogenic spectrum, displaying proportions of proliferating T, Treg, and NK cells that substantially surpassed all other cancer types, establishing a redefined immunogenicity hierarchy (laryngocarcinoma > skin > lung > breast) ^14,31^ (**Figure 4D**). mIHC validation confirmed that while LSCC and lung BrMs shared comparable overall T cell infiltration densities, LSCC lesions harbored a significantly higher fraction of Tregs (4.6% versus 1.29% of total T cells) and proliferating T cells (31% versus 16% of total T cells) (**Figure S4C**). These findings distinguish LSCC BrM as an ultra-hot metastatic subtype.

To resolve distinct T cell functional states, the compartment was sub-clustered into seven subsets: naïve, memory, cytotoxic CD8 T (CTL), exhausted CD8 T (Tex), interferon-responsive CD8 T, stress-state, and resting T cells (**Figures 4E and 4F**). In alignment with global lymphoid distribution, melanoma BrMs demonstrated the highest abundance across nearly all functional subsets, while breast BrMs consistently exhibited the lowest (**Figure 4G**). Deep functional and metabolic profiling targeted the core effector and suppressive populations: CTLs, Tex cells, and Tregs (**Figure 4H**). While CTLs displayed peak activation scores, Tex cells maintained robust activation and interferon-response signatures despite their exhausted state. Metabolically, Tex cells were characterized by maximal oxidative phosphorylation, glycolysis, and fatty acid metabolism pathways, closely followed by Tregs (**Figure 4H**). Tregs simultaneously exhibited the highest NF-κB signaling and cytokine-receptor activity, underscoring their potent suppressive machinery within the metastatic niche (**Figure 4H**).

An analysis of the CTL-to-Tex ratio revealed a striking paradox between cold and hot intracranial tumors (**Figure 4I**). Typically cold microenvironments, such as primary gliomas and breast BrMs, were heavily dominated by unexhausted CTLs, whereas hot malignancies (melanoma and laryngocarcinoma BrMs) were dominated by Tex cells (**Figure 4I**). This topological shift was validated via CD8 IHC and CD8/PD-1/GZMB mIHC across an independent validation cohort (**Figures 4J**). These data delineate dual, origin-dependent tracks of immune escape across the brain tumor spectrum. In cold tumors, rigorous immune exclusion limits T cell influx^32^; the scarce resident CTLs do not show exhaustion markers and likely represent non-tumor-specific bystander T cells^33^. Conversely, hot microenvironments feature massive T cell infiltration that, upon chronic antigen confrontation, is driven into a terminally exhausted yet hyper-metabolic state^34^. This structural bifurcation emphasizes the necessity for precision immunotherapies tailored to the distinct immune baselines inherited from the primary site.

### Functional Partitioning of Stromal-Vascular Remodeling and Angiocrinitic Specialization in Gliomas and Brain Metastases

Single-cell transcriptomic profiling of 11,188 stromal and vascular cells resolved smooth muscle cells (SMC.MYH11), pericytes (PC.01.PDGFRB, PC.02.RGS5), fibroblasts (FB.01.PDGFRA, FB.02.COL1A1, FB.03.MMP14), and specialized endothelial subsets spanning arterial (Art.EFNB2), venous (Ven.01.TSHZ2, Ven.02.ACKR1), capillary (Cap.01.CAVIN2, Cap.02.RGCC), and sprouting tip endothelial cells (Tip.PLVAP) (**Figure 5A**). Comparative quantification revealed a distinct compositional bifurcation between primary and secondary brain malignancies. Secondary BrMs exhibited a prominent expansion of CAFs and Tip.PLVAP cells, a phenotype particularly pronounced in lung, skin (melanoma), and breast metastases (**Figures 5B, 5D, and 5F**). Structurally, the BrM niche was enriched for FB.02.COL1A1, SMC.MYH11, Ven.02.ACKR1, Tip.PLVAP, and Art.EFNB2, whereas gliomas were fundamentally biased toward a quiescent capillary-and pericyte-centric framework dominated by PC.02.RGS5, Cap.01.CAVIN2, and Cap.02.RGCC (**Figure 5D**). This divergence indicates that metastatic colonization precipitates a multi-lineage structural transformation involving active matrix deposition, venous-like remodeling, mural rearrangement, and tip-like activation, whereas primary gliomas preserve native neurovascular configurations.

**Figure 5.**
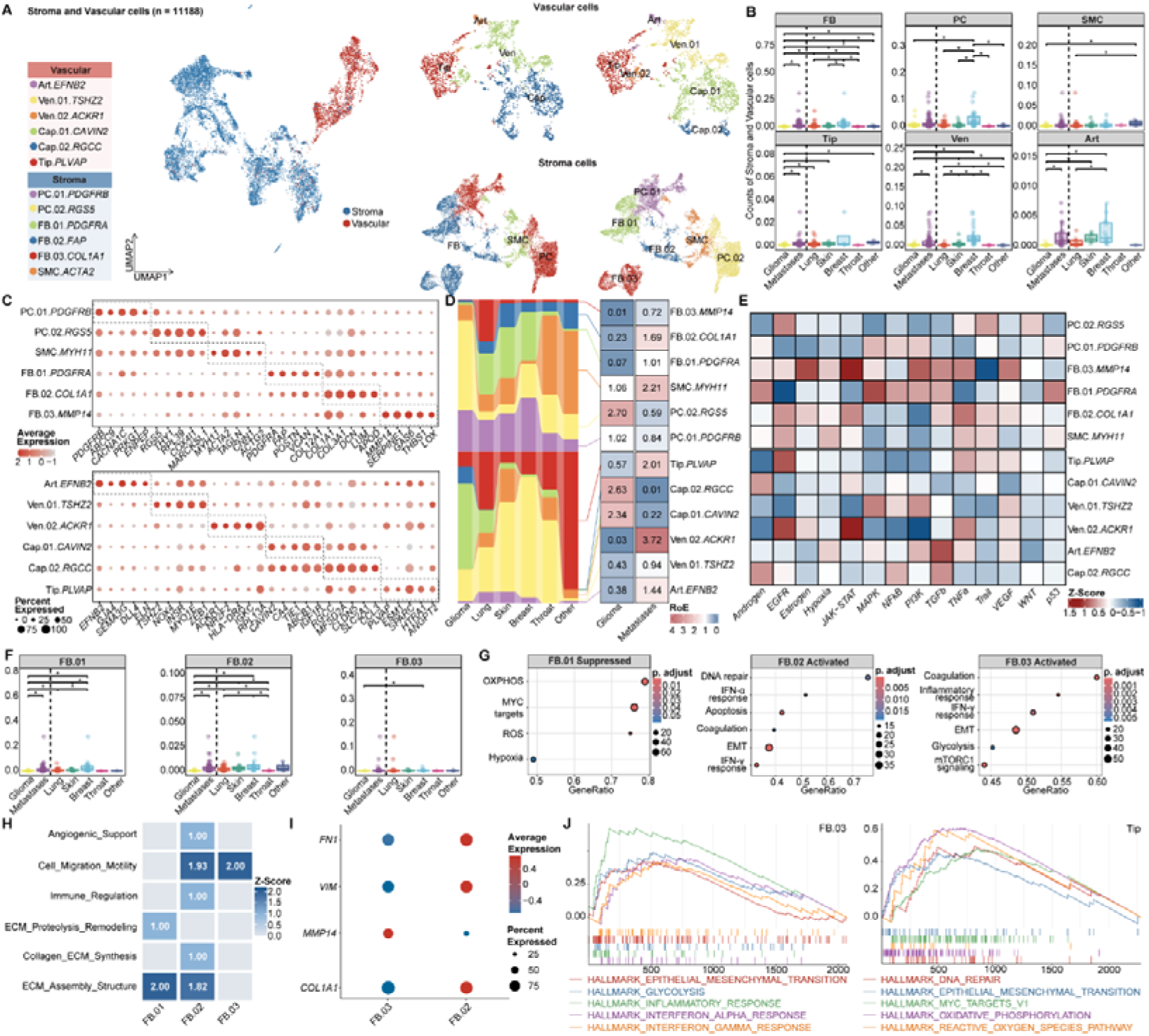
Single-cell Transcriptomic Dissection of Stromal and Vascular Remodeling Programs in Glioma and Brain Metastases. **(A)** UMAP embedding of 11,188 stromal and vascular cells annotated by high-resolution sub-clusters. **(B)** Box plots showing major stromal-vascular cluster abundance across cohorts. **(C)** Canonical marker profiles defining stromal and vascular sub-clusters. Significance evaluated by a two-sided Wilcoxon rank-sum test (asterisks denote adjusted P < 0.05). **(D)** Compositional distribution (Sankey) and matrix representation of observed-to-expected (ROE) enrichment across tumor types. **(E)** Scaled pathway activity heatmap across stromal and vascular sub-clusters. **(F)** Fibroblast sub-cluster abundance across cohorts. Significance evaluated by a two-sided Wilcoxon rank-sum test (asterisks denote adjusted P < 0.05). **(G–H)** Bubble plots showing pathway enrichment **(G)** and Z-score-normalized functional module scores **(H)** across fibroblast subsets and tip endothelial cells. **(I)** Comparative marker dot plot tracking epithelial-mesenchymal and proteolysis-related transitions between FB.03.MMP14 and FB.02.COL1A1. **(J)** GSEA curves showing activated functional pathways in FB.03.MMP14 and Tip.PLVAP.

Deconvolution of the CAF compartment revealed a functional duality between matrix deposition and proteolytic degradation. The FB.01.PDGFRA subset expanded preferentially in lung and skin BrMs, orchestrating ECM reorganization alongside the metabolic suppression of oxidative phosphorylation and MYC targets (**Figures 5C, 5F, 5G, and 5H**). The dominant FB.02.COL1A1 subset represented a matrix-deposition state overexpressing *COL1A1, FN1*, and *VIM*, exhibiting broad synthetic activity for collagenous scaffolds across multiple BrM origins (**Figures 5C, 5F, and 5H**). Crucially, the EMT signature in FB.02 cells was coupled with structural matrix construction rather than active proteolysis (**Figure 5G**). In contrast, the less abundant FB.03.MMP14 sub-cluster operated as a dedicated matrix-degradation engine characterized by *MMP14* expression, superior motility, and active matrix proteolysis (**Figures 5C, 5E, and 5G–5J**). Thus, CAFs split into a matrix-constructing arm (FB.02) that deposits structural scaffolding and an invasive matrix-degrading arm (FB.03) that carves out migratory routes. Internal heterogeneity within the FB.02 pool, including a tumor-suppressive *ISLR*-high fraction, suggests that indiscriminate pan-fibroblast ablation could eliminate host-protective stromal constraints^35^.

Within the vascular compartment, the Tip.PLVAP population emerged as an expanded, pro-angiogenic, and infiltrative endothelial state in BrMs, showing pronounced enrichment across lung, skin, and breast metastases (**Figures 5C, 5E, and 5F**). Specialized for accelerated migration, hyper-permeability, and active sprouting angiogenesis, Tip.PLVAP cells displayed transcriptomic activation of DNA repair, ROS metabolism, and EMT programs (**Figures 5D, 5G, and 5J**). Meta-gene evaluation confirmed a shared mesenchymal signature between Tip.PLVAP and FB.03.MMP14, though FB.03 was tailored toward proteolysis whereas Tip.PLVAP was dedicated to endothelial hyper-motility (**Figure S5C**). This endothelial specialization provides a cellular basis for BBB disruption, hyper-permeability, and vessel co-option, likely sustained via a coordinated paracrine circuit involving endothelial EGFR signaling, tumor-derived *ANGPT2*, and FB.03-secreted VEGF^36,37^.

Cross-cancer stratification demonstrated that while secondary BrMs undergo convergent microenvironmental remodeling, they retain unique, origin-specific preferences. Lung and skin BrMs exhibited a synchronized expansion of FB.01.PDGFRA and FB.02.COL1A1 with variable Tip.PLVAP activation, whereas breast BrMs favored a distinct blueprint dominated by FB.02.COL1A1 and Tip.PLVAP induction (**Figures 5B, 5D, and 5F**). The highly invasive FB.03.MMP14 state was restricted to localized infiltrative niches rather than manifesting as a global pan-cancer expansion, reflecting its specialized role in focal parenchymal seeding. Integrating these components into a unified framework reveals that distinct primary malignancies ultimately converge on shared stromal-vascular adaptation programs.

Collectively, these findings establish that stromal and vascular remodeling in secondary brain tumors represents an active microenvironmental adaptation rather than a passive byproduct of spatial expansion. CAF states strategically reshape physical boundaries through balanced matrix deposition and degradation, while tip-like endothelial cells drive aberrant neovascularization and barrier disruption, suggesting that effective therapeutics must target these shared microenvironmental dependencies alongside primary lineage markers.

### Spatial Profiling Defines a Functionally Partitioned Ecosystem of Mesenchymal-Like States in Brain Metastasis

Subcellular-resolution spatial transcriptomic profiling via CosMx SMI of 128,675 high-quality cells across 9 BrM specimens resolved seven major lineages: malignant cells, BMDMs, CAFs, vascular cells, astrocytes, B/plasma cells, and T/NK cells (**Figure 6A, upper panel, and Figures S6A–S6E**). Utilizing our single-cell reference atlas, malignant cells were partitioned into four conserved archetypes: Glandular, MES-like, Squamous/Mucinous, and Neuronal Mimicry (**Figures 6B–6D, and S6F–S6G**).

**Figure 6.**
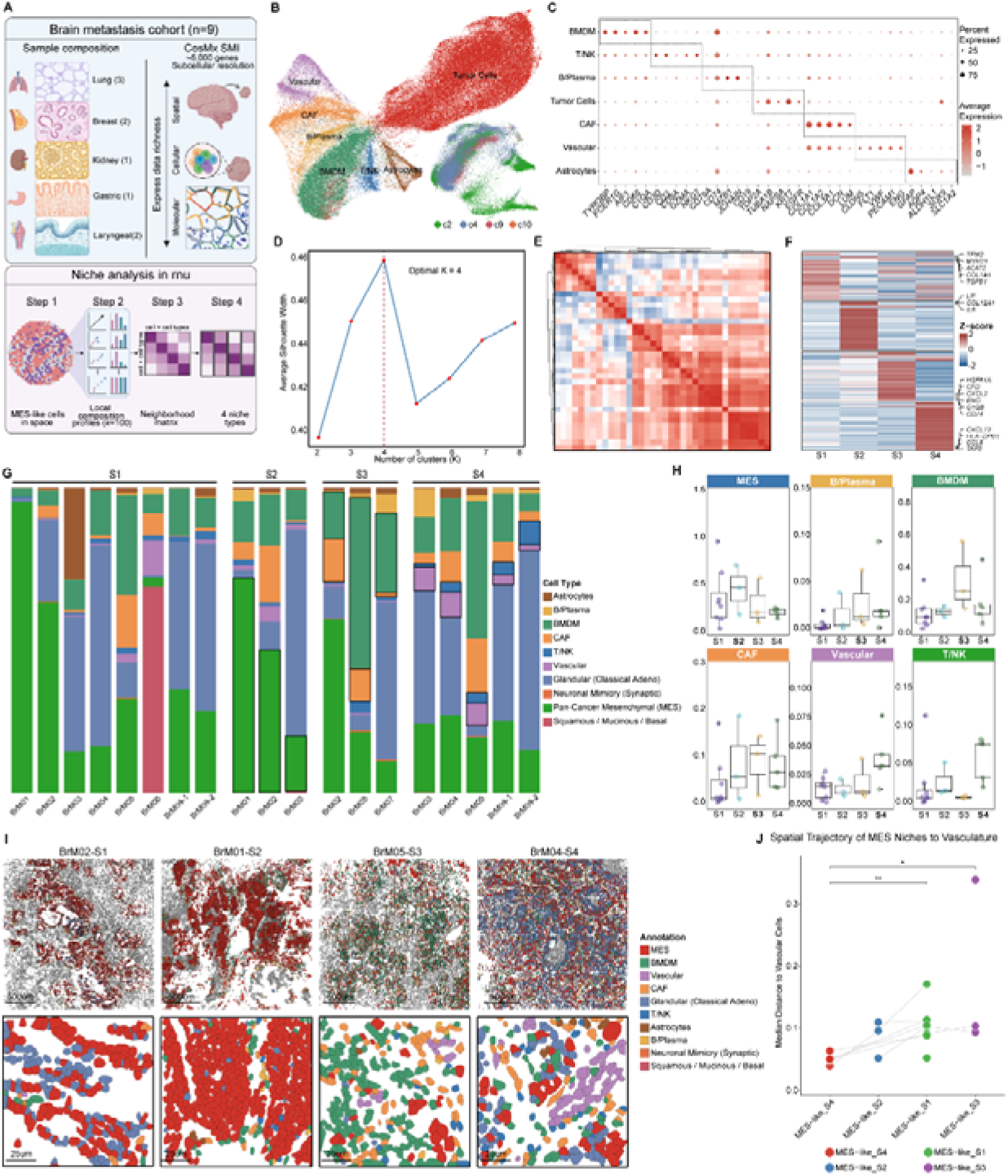
Spatial Profiling and Niche Architecture of Mesenchymal-Like States in Brain Metastases. **(A)** Graphical pipeline of CosMx 6k SMI spatial sample processing (top) and computational neighborhood deconvolution workflow for MES-like cells (bottom). **(B–C)** Global spatial UMAP of 128,675 cells annotated by major lineage **(B)** and malignant archetype reference mapping **(C)**. **(D)** Dual-layer marker dot plot for major lineages and malignant archetypes. **(E)** Niche partition optimization curve showing average silhouette width across K cluster dimensions. **(F)** Pairwise Pearson correlation heatmap of cellular neighborhood spatial niches based on local composition. **(G)** Differentially expressed signature genes across spatial niches S1 to S4. **(H–I)** Spatial niche compositional profiles showing distribution across tissue samples **(H)** and defining neighborhood cell frequencies **(I)**. **(J)** In situ cell-segmentation-resolution spatial layout maps of niches S1–S4 across representative sections, with 100-nearest neighbor contexts highlighted (top) and high-magnification fields included (bottom). **(K)** Tiered proximity model quantification measuring physical distance from index MES-like niche cells to functional vasculature across specimens.

To map the spatial architecture and local niches of the dominant, invasion-associated MES-like population, we employed a cellular neighborhood clustering strategy based on immediate spatial neighbors (**Figure 6A, lower panel**). This deconvolution identified four statistically robust and biologically distinct niche states (S1–S4) exhibiting striking functional divergence (**Figures 6E–6G**). The S1 (Homeostatic Core) state was enriched for drivers *YES1, TACSTD2*, and *ERBB2* alongside the stress-protective chaperone *CLU*, representing a stable, tumor-intrinsic core shielded from extrinsic perturbations. The S2 (Proliferative Hub) state overexpressed cell-cycle regulators (*STMN1, CENPF*) and stemness factors (*SOX2, MYB*), acting as rapid intralesional expansion centers. The S3 (Reactive Stroma) state was defined by a profound inflammatory and ECM remodeling signature, including *SPP1, CXCL8, CCL2*, and *COL4A1*. Finally, the S4 (Perivascular Sentinel) state exhibited robust antigen presentation (*CD74, HLA-DRA*), interferon-response genes (*IFI27*), and vascular stress markers (*SERPINE1, SAA1*), reflecting localized exposure to host circulatory and immune interfaces.

Quantification of neighbor compositions and spatial mapping onto tissue coordinates validated the topological segregation of these states (**Figures 6H–6J**). S1 niches demonstrated a simplified cellular background lacking peripheral immune or reactive stromal components, consistent with a sequestered tumor core localization. Conversely, S3 cells displayed a significant co-enrichment with BMDMs and CAFs. This spatial synergy establishes the MES-S3 state as a functional factory that recruits and educates myeloid cells via the *SPP1-CXCL8* axis, transforming the tumor margin into an immunosuppressive reactive stroma. The S4 state showed peak enrichment of vascular endothelial cells and T/NK cells, confirming its position as the frontline of hematogenous entry where immune surveillance and microvascular proliferation coexist^18^. In contrast, the S2 state was characterized by tight, homotypic aggregation of MES cells, which provides the physical constraints necessary for efficient clonal expansion while avoiding intermixture with other microenvironmental components.

Finally, physical distance quantification between MES cells and the nearest functional vasculature established a tiered proximity model (**Figure 6K**). S4 cells were localized at the immediate perivascular interface, while S2 cells were also physically proximal to vessels to satisfy the high metabolic demands of proliferative expansion. In contrast, both the S1 core and S3 reactive margin were significantly sequestered from functional vessels. The distal, vessel-depleted localization of S3 likely creates a localized hypoxic environment that drives the compensatory expression of chemoattractants (*SPP1* and *CXCL8*).

Together, these findings delineate a spatially stratified model of BrM evolution: MES-like cells initiate from a perivascular immune-adaptive outpost (S4), undergo rapid expansion in metabolically enriched proximal zones (S2), and ultimately diverge into either a homeostatic, sequestered core (S1) or a reactive, stroma-remodeling center at the invasive periphery (S3).

### Clinical Validation of MES-Like Spatial Niches Identifies Molecular Subtypes with Divergent Proteogenomic Landscapes and Prognostic Outcomes

To evaluate the translational potential of the identified MES-like spatial architectures, we projected the S1–S4 signatures onto a clinical cohort comprising transcriptomic and proteomic data from 312 BrMs. Unsupervised hierarchical clustering based on niche signature activity robustly partitioned patients into four distinct clinical subtypes: Proliferative, Angiogenic, Stroma-Reactive, and Quiescent (**Figure S7A**). Heatmap analysis revealed a direct topological correspondence between bulk clinical subtypes and single-cell spatial MES states: the Proliferative subtype enriched for the MES-S2 (proliferative expansion) signature, the Angiogenic subtype mapped to S4 (perivascular sentinel), the Stroma-Reactive subtype aligned with S3 (matrix remodeling), and the Quiescent subtype matched S1 (homeostatic core).

These niche-defined molecular subtypes exhibited significant prognostic divergence. Kaplan-Meier analysis of the transcriptomic cohort (**Figure S7B**; p = 0.0083) revealed that patients within the Proliferative group experienced the worst clinical outcomes and the shortest median survival, demonstrating that MES-S2-driven rapid intralesional expansion is a primary determinant of clinical deterioration. Parallel validation using an independent proteomic dataset (n = 107) confirmed the molecular stability of these niche states at the protein level (**Figure S7C**). However, proteomic survival analysis (**Figure S7D**; p = 0.0049) unveiled a transcript-protein decoupling phenomenon: while the Proliferative subtype maintained its poor prognosis, the Quiescent subtype also exhibited compromised survival at the protein level. This discordance suggests that although S1-state cells in the homeostatic core remain transcriptionally quiescent, they may accumulate superior stress-resistance factors or stable protein networks that confer late-stage malignancy. Conversely, the Angiogenic and Stroma-Reactive subtypes demonstrated relatively favorable survival profiles across both omics layers.

Characterization of clinical metadata revealed distinct lineage-specific constraints among these spatial niches (**Figures 7E and 7F**). Lung cancer BrMs were significantly enriched within the Angiogenic subtype, indicating a strong biological dependency on the perivascular niche, whereas breast cancer BrMs preferentially accumulated within the Proliferative subtype. These data confirm that while secondary brain malignancies undergo convergent microenvironmental evolution, their specific spatial survival strategies remain restricted by their primary tissue of origin.

Finally, cross-referencing our spatial framework against established bulk BrMS molecular subtypes demonstrated that the Proliferative subtype closely aligns with the poor-prognosis BrMS4 (proliferative) cluster, whereas the Angiogenic subtype is tightly linked with the immune-hot BrMS2 (immune-infiltrated) classification (**Figure S7G**). Consistent with previous reference archetypes, this BrMS4/Proliferative subtype is defined by high CIN, frequent *RB1* mutations, and APOBEC-associated mutational signatures that drive cell cycle dysregulation^38^.

Collectively, these findings indicate that the MES-like spatial niche framework provides a powerful tool for clinical risk stratification across proteogenomic dimensions, offering a cellular-dynamics-based explanation for bulk omics heterogeneity and establishing a theoretical foundation for niche-targeted precision interventions.

## Discussion

The brain, an “evolutionary island” characterized by its physiological isolation and stringent metabolic selectivity, imposes a unique selective pressure that shapes the intra-tumoral heterogeneity of resident malignancies. In this study, we constructed a large-scale single-cell transcriptomic atlas comprising 554,679 cells, further augmented by high-resolution spatial transcriptomics and clinical proteomics. By performing a cross-lineage comparison between gliomas and a diverse cohort of BrM, we transcended static cellular indexing to define the systemic logic of how malignant cells breach the blood-brain barrier and adapt to the neural niche. This multi-modal integration shifts our understanding of brain-resident tumors from a descriptive census of cell types to a dynamic model of spatiotemporal evolution and clinical relevance.

The phenotypic evolution of malignant cells upon colonizing the brain represents a profound tug-of-war between ancestral “lineage imprinting” and the acute selective pressures of the local microenvironment. Our findings demonstrate that while BrMs of diverse origins retain distinct signatures of their primary tissues, the brain environment drives a robust program of convergent evolution. Specifically, the mesenchymal-like state (MES-like, c2 program) emerges as a central functional hub. This state is not only intrinsically linked to tumor invasiveness but also serves as a shared conduit for immune evasion and therapeutic resistance. Concurrently, the identification of a “neuronal mimicry” state (c10 program), characterized by the expression of synaptic genes such as *SYT1*, underscores an extreme adaptive strategy where tumor cells functionally integrate into host neural circuits. These observations suggest that heterogeneous malignant cells converge upon a “universal survival grammar” to thrive in the neural niche, with the MES-like state acting as the core functional verb driving this adaptation.

As malignant cells evolve, the TME undergoes a dramatic metamorphosis, transitioning from an architecture dominated by brain-resident glia to one remodeled by peripheral infiltrates. In contrast to the microglia-centric landscape of gliomas, BrMs are characterized by a massive influx of bone marrow-derived macrophages and neutrophils, creating a more antagonistic immunological arena. Although BrMs often exhibit higher T-cell infiltration compared to gliomas, we unveil a profound “immune paradox”: in “hot” BrMs such as melanoma, the abundant T cells exist in a state of terminal exhaustion and hyper-metabolic dysfunction, resulting in a functional standoff where the immune system is present but paralyzed. This TME reorganization, orchestrated by the functional division of specialized vascular endothelia (Tip.PLVAP) and fibroblasts (FB.02/03), provides the essential physical scaffold and immunological shield for the convergent evolution of malignant cells.

A key innovation of this study is the definition of spatial niches (S1–S4) via spatial transcriptomics, which assigns biological meaning to the topological architecture of the tumor. By coupling these niches with clinical proteomic cohorts, we mapped spatial distribution directly onto clinical outcomes. The S1–S4 framework provides a chronological model of brain colonization: malignant cells initiate invasion at the perivascular “sentinel” niche (S4), enter a burst of proliferation in the nutrient-rich proximal vascular niche (S2), and subsequently diversify into the stromal-remodeled periphery (S3) or the metabolic core (S1). Crucially, this evolutionarily-derived spatial logic was validated in clinical cohorts, where a high S2 (proliferative) signature significantly correlated with the poorest patient survival. This suggests that the spatial topology of brain tumors is not a stochastic assembly of cells but an organized ecosystem driven by a temporal logic of colonization that ultimately dictates patient prognosis.

### Limitation

Despite the comprehensive nature of our multi-omic and spatial atlas, several limitations warrant consideration. First, while our cohort covers major primary origins, certain rare types (e.g., laryngocarcinoma) are represented by a limited number of cases, necessitating caution when generalizing these findings to broader populations. Second, our spatial models are derived from cross-sectional surgical resections of established lesions; future studies utilizing longitudinal samples or animal models are essential to capture the earliest stages of metastatic seeding and dynamic evolution under therapeutic pressure. Third, given the limited sample size of our spatial cohort, this dimension of the study represents a pilot exploration rather than a comprehensive spatial atlas. Additionally, the platform relies on a predefined targeted gene panel (6k) rather than unbiased whole-transcriptome sequencing, potentially omitting unselected regulatory transcripts or non-coding elements. Fourth, projecting single-cell spatial niche signatures onto bulk clinical proteogenomic cohorts relies on computational deconvolution, which inherently lacks direct *in situ* validation of topological boundaries in large retrospective series. Finally, while we identified SYT1 and the MES-like states as pivotal, the precise molecular switches governing these phenotypic transitions and their viability as therapeutic targets require rigorous functional validation.

## Methods

### Patients and Sample Collection

Patients were prospectively enrolled into this study at First Affiliated Hospital of Soochow University. Written informed consent was provided by all participants according to approved protocol. Fresh tumor tissue was collected at the time of surgery after confirmation of metastasis on frozen section. Patients who had received any prior local or systemic therapies—specifically radiotherapy (e.g., Stereotactic Radiosurgery or Whole-Brain Radiotherapy), or systemic treatments including chemotherapy, targeted therapy, or immunotherapy within one month prior to neurosurgical resection—were excluded from the study. Tumors were categorized according to the 2021 WHO Classification of Central Nervous System Tumors based on the combination of relevant histopathologic and molecular features from IHC.

### Single cell RNA library preparation and sequencing

Patient tumors were mechanically cut into about 1-2 mm3 tissue in Dulbecco’s modified eagle medium (DMEM) (Gibco). They were then enzymatically digested with the human tumor dissociation kit (Miltenyi Biotec) on the gentleMACS Dissociator (Miltenyi Biotec) per manufacturer instructions. Dissociated cells were filtered by a 70-mm strainer and centrifuged at 300g for 7 min. After removing the supernatant, the pelleted cells were suspended in red blood cell lysis buffer (Miltenyi Biotec) for 2 min to remove red blood cells. The cells were then washed with sorting buffer [phosphate-buffered saline (PBS) supplemented with 2% fetal bovine serum (FBS)] and accessed for viability using trypan blue exclusion. 10,000 - 20,000 cells were loaded for library construction using the 10x Chromium Single cell 5‘Library (10x Genomics, V3), according to the manufacturer’s instructions. Purified libraries were subject to an Hiseq X Ten sequencer (Illumina) for sequencing with 150-bp paired-end reads.

### Single-cell RNA-seq data preprocessing

Newly generated single-cell sequencing data were aligned with the GRCh38 human reference genome and quantified using Cell Ranger (version 3.0, 10x Genomics). The preliminary filtered data generated from Cell Ranger were used for downstream filtering and analyses. The quality of cells was then assessed based on two metrics: (1) The number of detected genes per cell; (2) The proportion of mitochondrial gene (Genes starting with ‘MT-’) counts per cell; (3) The proportion of ribosomal gene (genes starting with ‘RPS’ or ‘RPL’) counts per cell; (4) The proportion of hemoglobin gene (genes starting with ‘HB’ but are not followed by ‘P’) counts per cell. Specifically, cells with detected genes fewer than 500 or more than 5000, mitochondrial unique molecular identifier (UMI) count percentage larger than 10%, lysosomal UMI count percentage larger than 50%, and hemoglobin UMI count percentage larger than 0.05% were filtered out. To remove the potential doublets, Scrublet^39^ was used for each sequencing library with the expected doublet rate set to be 0.05, and cells with the predicted doublet Score larger than 0.3 were further filtered out. Finally, we excluded samples with fewer than 500 retained cells. For scRNA-seq data from other publications, the same filtering steps were applied for the datasets with raw count matrix to obtain high-quality cells. The detailed metadata (including tissue location, sample and patient identifier) were retrieved from the original studies (Table S1).

### Batch effect correction

We next normalized each strictly filtered count data using the *scanpy.normalize_total* function with parameter ‘*target_sum=1e4*’ in Scanpy^40^ (v1.10.1). All the normalized data were logarithmically transformed for downstream analyses. The unexpected effects of the total counts and the original identify were regressed out from the normalized expression matrix using the *scanpy.pp.regress_out* function with the parameter setting. Batch effects between samples were preliminarily corrected by simple linear regression. We then used *scanpy.concatenate* function to merge the dataset processed by the unified process together at the data structure level with the parameter ‘*join = ‘inner*’’. Batch effects between datasets were preliminarily corrected by merging standardized datasets rather than directly merging count matrices.

For the merged matrices, we scaled each gene to unit variance using sc.pp.scale function with the parameter ‘max_value=10’, and then highly variable genes (HVGs) were selected using the *scanpy.pp.highly_variable_genes* function with default parameters. Next, principal component analysis (PCA) was performed on the matrix of HVGs to reduce noise and reveal the main axes of variation using the *scanpy.tl.pca* function, and the top 30 components were retained for downstream analyses. The batch effects were corrected by the BBKNN^41^ algorithm (v1.6.0), which detected the top nearest neighbors of each cell from each batch respectively instead of the entire cell pool. The parameters of the *scanpy.external.pp.bbknn* function were set to ‘‘*batch_key=’dataset’, n_pcs=30*’’. As comparison of retaining batch effects, the neighborhood graph was computed using *scanpy.pp.neighbors* function with default parameters following PCA. Finally, Uniform Manifold Approximation and Projection (UMAP) was employed for visualization via the *scanpy.tl.umap* function with default parameters.

### Preliminary annotation

We performed the unsupervised clustering to unveil the structure of brain malignancy population using the *scanpy.tl.leiden* function with the parameter ‘*resolution=2.0*’ and the first round annotation was performed based on the expression of a predefined dictionary of canonical markers, including myeloid cells (*CD68, C1QA*), lymphocytes (*CD3D, PTPRC*), vascular (endothelial) cells (*VWF, PECAM1, ENG*), and stromal cells (*COL1A1, COL1A2, LUM*). Malignant cells were characterized by the expression of *PTPRZ1, BCAN, SOX2, OLIG2, EGFR*, as well as primary-specific markers such as *NKX2-1, SFTPC, NAPSA* (lung-derived) and *KRT7, KRT18, KRT19* (epithelial-derived). Finally, oligodendrocytes were defined by *PLP1, CNP, CLDN11, MBP,* and *MOBP*. The cluster-specific marker genes were identified using the *scanpy.tl.rank_genes_groups* function with the parameter of ‘*method=t-test*’.

### Further annotation

For Malignant cells, we performed unsupervised clustering of malignant cells using cNMF implemented in Omicverse^42^ (v1.6.10). The malignant cell count matrix was first preprocessed using *omicverse.pp.preprocess* with the parameter “mode=’shiftlog|pearson’, n_HVGs=2000” to select highly variable genes, followed by scaling (*omicverse.pp.scale*) and principal component analysis (*omicverse.pp.pca*). We then initialized cNMF decomposition using *omicverse.single.cNMF* function with the parameter ‘*components=np.arange(2,20), n_iter=20, num_highvar_genes=2000*’, performing factorization using *omicverse.factorize* function before combining results using *omicverse.combine* function, both with default parameter. To find the optimal number of programs, the *k_selection_plot* function with default parameter was used to calculate the stability and reconstruction error. Finally, consensus clustering was performed using the *omicverse.consensus* function with the parameter ‘k=15, density_threshold=0.2’. Programs showing marked overexpression of myeloid, lymphoid, endothelial, stromal, or oligodendrocyte markers and containing relatively few cells were identified as contamination and subsequently excluded from further analysis.

The characterization of non-malignant cells proceeded through sequential classification phases, beginning with lineage assignment using canonical markers: myeloid populations included microglia-derived macrophages (MDMs; *TMEM119, P2RY12, CX3CR1*), bone marrow-derived macrophages (BMDMs; *CD14, FCGR3A/CD16, CD163, MRC1/CD206*), neutrophils (*S100A8, S100A9, LYZ*), dendritic cells (DCs, *CCR7, CLEC9A, FCER1A*) and mast cells (*CPA3, TPSAB1*); lymphoid populations comprised T cells (*CD4, CD8A, GZMK, LAG3, PDCD1*) including regulatory T cells (*FOXP3, CTLA4, TIGIT, IL2RA, TNFRSF18*), B cells (*CD79A, MS4A1/CD20*), plasma cells (*JCHAIN, XBP1, SDC1/CD138*), and natural killer cells (NKs, *NKG7, NCAM1/CD56, FCGR3A/CD16, KLRC1, FGFBP2*); endothelial cells (ECs) were subclassified as arterial (*EFNB2, GJA4, SEMA3G*), venous (*NR2F2, ACKR1*), capillary (*RGCC, CA4*), and tip-like (*PLVAP, ANGPT2*) variants; while stromal components included smooth muscle cells (SMCs, *ACTA2, TAGLN*), pericytes (PCs, *PDGFRB, CSPG4, RGS5*), and cancer associated fibroblasts (CAF, *COL1A1, COL3A1, DCN, PDGFRA, FAP*).

### Quantitative Analysis of Cell Cluster Distribution Preferences

Building upon the STARTRAC-dist framework developed by Zhang et al.^43^, we implemented a robust quantitative approach to assess cell cluster distribution patterns across treatment groups. Our analytical pipeline calculated the ratio of observed-to-expected cell frequencies (*R*_o/e_) to precisely quantify cluster-level enrichment or depletion according the following formula:

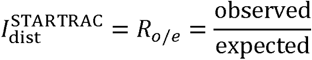

In which expected values were derived from chi-squared test contingency tables comparing cell clusters against treatment groups. This method provides distinct advantages over conventional statistical testing by: (1) generating continuous enrichment scores rather than binary significance thresholds. For example, if *R*_o/e_ > 1, it suggests that cells of the given cell cluster are more frequently observed than random expectations in the specific treatment group, that is, enriched. If *R*o/e < 1, it suggests that cells of the given cell cluster are less frequently observed than random expectations in the specific treatment group, that is, depleted; (2) maintaining directional information about cluster redistribution that is lost in traditional chi-squared values, which are defined as 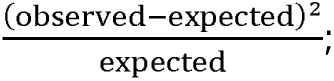 and (3) enabling cross-comparison of distribution preferences among all cell clusters within our annotated taxonomy.

### Jaccard Index

To cross-compare the transcriptomic similarity of subsets of microglial-derived macrophages between the Glioma and BrM, a Jaccard similarity index analysis was employed. From the integrated dataset, a representative subset of 10,000 cells was randomly sampled. Celltype specific marker genes were identified using the Seurat FindMarkers function with the parameter. The top 100 highly enriched markers for each cell type, ranked by decreasing average log2 fold change, were selected as corresponding transcriptomic signatures. The pairwise overlap between Glioma and BrM clusters was quantified by the Jaccard similarity index (J), calculated as:

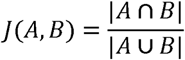

where A and B denote the top 100 marker gene sets of the respective clusters. The resulting similarity matrix was visualized as a non-clustered heatmap, with an asterisk (*) highlighting cell type pairs with strong transcriptional correspondence (J ≥ 0.3).

### Chromosomal Instability Assessment

To characterize genomic instability at single-cell resolution, we inferred copy number variations (CNVs) and calculated cumulative CNV scores for non-immune and non-vascular cells. Initially, inferCNV (v1.22.0)^44^ was employed to estimate CNV profiles based on single-cell transcriptomic data. Oligodendrocytes were designated as the reference diploid population to establish a baseline. The inference was performed using the *run* function with the parameter “utoff = 0.1, denoise = TRUE”. To quantify the overall magnitude of chromosomal alterations, we calculated the Mean Squared Deviation (MSD) for each cell, defined as the average of the squared transition values across all genomic windows. A higher MSD value represents a greater degree of global CNV and higher genomic instability. To ensure the robustness of our findings, we independently validated the CNV results using fastCNV (v1.1.1)^45^. Genomic scores were generated with a diploid baseline of zero. Similarly, the MSD for each cell was calculated as the average squared deviation of all genes from the baseline to represent the chromosomal instability (CIN) level. The high concordance between inferCNV and fastCNV results confirmed the reliability of our single-cell genomic instability quantification.

### Developmental Potential Quantification Using CytoTRACE 2

To assess cellular developmental potential across our single-cell atlas, we employed CytoTRACE 2^46^ (v1.1.0), an interpretable deep learning framework that predicts absolute developmental potential from scRNA-seq data. This validated algorithm classifies cells into discrete potency categories (totipotent, pluripotent, multipotent, oligopotent, unipotent, and differentiated states) while simultaneously generating continuous developmental potential scores ranging from 0 (terminally differentiated) to 1 (totipotent). We applied the *cytotrace2()* function with default parameters to leverage the algorithm’s deep neural network architecture trained on diverse developmental datasets to infer developmental potential based on gene expression patterns, where higher CytoTRACE scores (approaching 1) indicate greater developmental plasticity and lower scores reflect lineage commitment.

### Cell Type Benchmarking with Published Signatures

To systematically annotate and map cellular states to defined functional phenotypes across major lineages, we benchmarked cluster-specific transcriptomic profiles against established reference frameworks utilizing the Seurat *AddModuleScore* pipeline. For each distinct subpopulation within the malignant, fibroblast, neutrophil, and broader myeloid compartments, lineage-specific signatures were initially derived by extracting the top 100 DEGs via the *FindMarkers* function.

To resolve malignant cell heterogeneity, enrichment scoring was executed across three orthogonal axis frameworks, integrating (1) 41 consensus pan-cancer meta-programs (MPs) from Gavish et al^47^, (2) 8 brain metastasis-specific functional MPs (including cell cycle, stress, and EMT modules) defined by Gonzalez et al^10^, and (3) 19 brain-adaptive modules (encompassing specialized neural-like states) characterized by Xing et al^5^. For the non-malignant microenvironmental architecture, granular activation states and functional trajectories of the overarching myeloid and neutrophil compartments were comprehensively mapped by evaluating single-cell profiling scores against (1) 20 brain-metastasis-associated myeloid cell programs from Xing et al^5^ and (2) 10 transcriptionally distinct states defining the neutrophil architectural framework established by Daniela et al^48^. Concurrently, functional specialization and phenotypic states within the lymphoid compartment were deconvoluted by scoring T cell profiles against the pan-cancer T cell atlas framework from Chu et al^49^, assessing four multidimensional regulatory modules as structured in 3 differentiation states, 10 functional signaling pathways, 3 metabolic programs and 2 apoptotic axes. In parallel, CAFs states were cross-referenced against three distinct stromal reference matrices, assessing alignment with (1) 20 cross-tissue fibroblast programs from Yang et al^50^, (2) 5 brain metastasis-associated CAF programs from Xing et al.^5^, and (3) 2 CAF or mesenchymal stem cell (MSC)-like programs from Gonzalez et al^10^.

### Pathway Enrichment analysis

Pathway enrichment was assessed using two complementary computational frameworks to comprehensively characterize biological processes across cell subpopulations. First, single-cell pathway activity was inferred via the multivariate linear model (MLM) implemented in the decoupleR package (v2.9.7)^51^. This algorithm, executed via the *run_mlm* function with the parameters “.source =’source’,.target =’target’,.mor =’weight’, minsize = 5”, was applied to the log-normalized expression matrix to evaluate two distinct regulatory networks. The canonical network utilized the PROGENy model to quantify 14 major signaling pathways (Androgen, EGFR, Estrogen, Hypoxia, JAK-STAT, MAPK, NFkB, p53, PI3K, TGFb, TNFa, Trail, VEGF, and WNT) based on the top 500 responsive genes ranked by P-value. Concurrently, to specifically delineate tumor-associated processes, we evaluated 10 manually curated functional signatures (IA, TPI, GIM, SPS, AID, ERI, RCD, EGS, DCE, and AIM). Each custom signature comprised hallmark genes assigned uniform positive regulatory weights (mor = 1). Gene set enrichment analysis (GSEA) was performed on specific cell subpopulations. For each subpopulation, differentially expressed genes were first identified using the *FindMarkers* function in Seurat (v4.3.0), followed by selection of the top 100 most significant genes based on statistical metrics. These gene sets were then analyzed using the *GSEA()* function from clusterProfiler^52^ (v4.14.6) package with two distinct collections from the Molecular Signatures Database (MSigDB): HALLMARK gene sets (c1.hallmark) and the legacy KEGG pathway collection (c2_kegg_legacy).

### Gene Regulatory Network Inference and Transcription Factor Activity Analysis

Gene regulatory network (GRN) inference was performed using the pySCENIC pipeline^53^ (v0.12.1). To address data sparsity, we constructed metacells for each subpopulation. Specifically, we performed nearest-neighbor graph construction using the *FindNeighbors* function with parameter “k.param=10”, followed by high-resolution clustering using the *FindClusters* function with parameter “resolution=50” to identify metacell groupings. Metacell expression profiles were generated by summing raw counts across all cells within each cluster. Metacells containing fewer than 10 cells were filtered out. The aggregated metacell matrix was further filtered to retain genes expressed in at least 5 metacells and genes present in the hg38 cisTarget database. GRN inference was performed using GRNBoost2 (arboreto_with_multiprocessing.py) and a human TF list derived from motifs-v10nr_clust-nr.hgnc-m0.001-o0.0.tbl. Putative regulons were refined using cis-regulatory motif analysis (pyscenic ctx) against both promoter (hg38_500bp_up_100bp_down_full_tx_v10_clust.genes_vs_motifs.rankings.feather) and extended genomic regions (hg38_10kbp_up_10kbp_down_full_tx_v10_clust.genes_vs_motifs.rankings.feather) databases. Regulon activity scores were calculated for individual cells using the AUCell algorithm implemented via the *ComputeModuleScore* function in Seurat (v4.3.0), with results stored as a new assay for downstream analysis.

### Integration and Reference Mapping of an External breast cancer BrM Cohort

To correct the lineage imbalance of breast cancer BrM in our master atlas (initially n = 4 versus lung cancer BrM n = 59 and melanoma n = 64), we integrated an external large-scale breast cancer BrM^54^. Raw sparse matrices were partitioned into Malignant, Myeloid, Lymphoid, Endothelial, and Stromal compartments and initialized as Seurat objects. To preserve our established batch-corrected framework, external cells were projected onto the master reference using a supervised reference-mapping workflow rather than global re-embedding. Briefly, the annotated master reference was log-normalized, subsetted for the top 2,000 highly variable genes, and scaled for PCA and UMAP modeling. Query compartments underwent identical preprocessing. Transfer anchors were then computed via *FindTransferAnchors* function with parameter “reference.reduction =’pca’, dims = 1:50”, and fine-grained cell-type identities were projected onto the query cells using *TransferData* function. Transferred annotations, metadata, and expression profiles were fully integrated into all downstream statistical meta-analyses, compositional profiling, and cross-lineage characterizations, though these external cells were excluded from the primary master UMAP coordinates.

### Immunohistochemistry (IHC)

Formalin-fixed paraffin-embedded (FFPE) sections were subjected to IHC staining to evaluate **CD8** expression (Proteintech, 66868-1-Ig). Briefly, sections were deparaffinized in xylene and rehydrated through a graded series of ethanol. Antigen retrieval was performed in a microwave-heated retrieval buffer at high power until boiling, followed by maintenance at low heat for 15 min. To eliminate endogenous peroxidase activity, sections were treated with 3% hydrogen peroxide for 10 min in the dark. After blocking with goat serum at room temperature for 30 min, slides were incubated with the primary anti-CD8 antibody for 1 hour in a humidified chamber. Subsequently, sections were incubated with an HRP-conjugated secondary antibody for 10 min, followed by visualization using a DAB (3,3’-diaminobenzidine) substrate. Sections were then counterstained with hematoxylin, differentiated with 1% acid ethanol, and blued under running water. Finally, the slides were dehydrated, cleared in xylene, and mounted with neutral resin. All stained sections were digitized using a high-resolution slide scanner for further analysis.

### Multiplex Immunohistochemistry (mIHC)

mIHC was performed using a sequential staining strategy to characterize the glioma/BrM microenvironment. Briefly, 4-μm-thick FFPE sections were deparaffinized in Xylene, rehydrated through a graded series of ethanol (100%, 90%, and 70%), and rinsed in distilled water. Antigen retrieval was conducted via microwave heating in a dedicated retrieval solution at high power until boiling, followed by 15 min at low power. Endogenous peroxidase activity was quenched with 3% H_2_O for 10 min, and non-specific binding was blocked with normal goat serum for 30 min at room temperature. The sections were then subjected to multiple rounds of sequential staining. In each round, slides were incubated with primary antibodies for 1 h, followed by secondary antibody incubation (10 min) and signal amplification using tyramide signal amplification (TSA)-based fluorescent dyes (10 min). To enable multi-target labeling, the previous primary-secondary antibody complexes were stripped between each round using microwave-based heat treatment. Three specialized panels were employed: Panel 1 (Tumor/Neuron) included SYT1 (YT4484, ImmunoWay; Opal 620) and Pan-CK (ZM-0069, Zsbio Tech; Opal 520); Panel 2 (Immune/Proliferation) included CD56 (14255-1-AP, Proteintech; Opal 620), Foxp3 (ab20034, Abcam; Opal 570), CD3 (ab16669, Abcam; Opal 520), and Ki67 (ab16667, Abcam; Opal 690); and Panel 3 (Effector/Exhaustion) included GZMB (ab255598, Abcam; Opal 620), PD-1 (ZM-0381, Zsbio Tech; Opal 520), and CD8 (66868-1-Ig, Proteintech; Opal 690). Following the final staining round, nuclei were counterstained with DAPI for 10 min. Slides were then mounted with fluorescence mounting medium and scanned using a high-resolution slide scanner to generate multichannel images.

### CosMx Spatial Transcriptomic Profiling and Cell Segmentation

Spatial transcriptomic profiling was performed on 5μm FFPE sections from nine samples (obtained from eight patients) using the CosMx Spatial Molecular Imager (SMI) (NanoString Technologies) equipped with the 6000-plex Human Universal Cell Characterization Panel. The experimental workflow followed a rigorous multi-day protocol. Briefly, FFPE sections were dried at 65°C, deparaffinized in xylene, and rehydrated through a graded ethanol series. Heat-induced antigen retrieval was executed at 100°C for 15 min using a pressure cooker (BioSB), followed by stabilization in DEPC-treated water. Tissues were then permeabilized with a specialized digestion buffer at 40°C for 30 min. To facilitate precise image registration, fiducial markers (0.001%) were applied and incubated for 5 min. Following post-fixation in 10% neutral buffered formalin (NBF), sections were treated with Sulfo-NHS-acetate for 15 min to reduce background autofluorescence. For in situ hybridization, the CosMx RNA probe mixture was denatured at 95°C for 2 min and hybridized to the sections at 37°C overnight.

On the second day, stringency washes were performed twice for 25 min each using a pre-warmed 50% formamide/4×SSC solution at 37°C. To ensure high-fidelity cell boundary detection within the complex brain microenvironment, sections were incubated for 1 hour with the CosMx Human Neuroscience Cell Segmentation Kit. This specialized cocktail enabled a four-channel morphology-based segmentation strategy: DAPI for nuclei (Ch1), Hs Neuro rRNA for neuronal cytoplasm (Ch2), Mm/Hs Neuro Histone for nuclear proteins (Ch3), and Mm/Hs GFAP for glial cytoplasm (Ch4). Finally, the slides were assembled into flow cells, and cyclic readout was initiated on the CosMx instrument. Raw data were processed via the AtoMx Spatial Informatics Platform for transcript decoding and cell-level assignment, yielding a single-cell spatial expression matrix. For quality control, cells with fewer than 100 detected transcripts (nFeature_RNA < 100) were excluded from downstream analysis.

### Niche Analysis of MES-like Cells via CosMx SMI

To define the localized cellular architecture surrounding MES-like cells, we conducted spatial niche analysis mainly using Seurat package (v5.1.0), FNN package^55^ (v1.1.4.1) and RcppML^56^ package (v0.3.7). For each individual MES-like cell, its spatial neighborhood was captured by identifying its 100-nearest neighbors (k = 100) based on 2D spatial coordinates via the *get.knnx* function in FNN package. A localized composition matrix was constructed by calculating the proportional frequency of 11 major cell types within this k=100 window. To decompose these complex localized environments into recurrent spatial patterns, NMF was performed on the normalized niche matrix using *nmf* function with the parameters “k = 5” in RcppML package. The resulting NMF embeddings (W-matrix) were integrated into the Seurat object as a dimensional reduction and utilized for unsupervised sub-clustering of MES-like cells within each sample using *FindNeighbors* function with parameter “reduction =’nmf’, dims = 1:5” and *FindClusters* function with parameter “resolution = 0.1, algorithm = 1” in Seurat package.

To enable a robust comparison across the nine samples, we integrated the niche sub-clusters while implementing a systematic background correction strategy. Recognizing that local cell-type enrichment can be confounded by tissue-wide abundance, we calculated the global cell-type fractions for each whole-tissue section. The average niche composition of each sub-cluster was normalized by its corresponding tissue background to generate a corrected enrichment score. These scores were then re-normalized to a sum of 1 to represent the relative enrichment of cell types within each niche. To ensure balanced representation across the cohort, a stratified downsampling strategy was applied, retaining a representative number of niche sub-clusters per sample to prevent specimens with high cell density from dominating the global landscape.

Conserved spatial functional units, termed “Global MES Niches,” were identified by performing hierarchical clustering on the integrated and balanced niche correlation matrix. The optimal number of clusters (K) was objectively determined using the *silhouette* function in cluster package (v2.1.8.1), which maximized the average silhouette width across a range of K from 2 to 8. Global clustering was executed via the *hclust* function with parameter “method =’ward.D2’” parameter based on Pearson correlation coefficients (PCC) in stats package (v4.4.2). This hierarchical workflow partitioned the MES-like neighborhoods into four distinct global niches (S1–S4). Differential expression analysis was subsequently conducted using the *FindAllMarkers* function with default parameters to identify niche-specific molecular signatures.

### Niche-based Clinical Stratification and Prognostic Analysis

To translate spatial niche architectures into clinical phenotypes, optimized molecular signatures for the four MES-like states (S1–S4) were developed by intersecting spatial markers with bulk-level features. A dynamic feature-selection strategy was implemented to account for differential coverage between omics platforms, refining signatures to the top 50 genes for transcriptomics (n=314) and top 40 proteins for proteomics (n=107), thereby ensuring robust representation across clinical samples. Individual niche activity was quantified using *gsva* function in GSVA^57^ package to transform gene-level expression into functional niche-activity profiles. Based on these profiles, unsupervised hierarchical clustering was performed, with the cluster number pre-specified to four to align with the identified spatial niches—Proliferative, Angiogenic, Stroma-Reactive, and Quiescent. The resulting classification was subsequently evaluated for prognostic significance and evolutionary convergence across diverse primary tumor origins.

## QUANTIFICATION AND STATISTICAL ANALYSIS

All data were analyzed and processed using R (v4.3.2) and Python (v3.10.12), with details of specific functions and libraries provided in the Methods sections. Statistical significance was determined by Mann-Whitney U test (Wilcoxon rank-sum test), Fisher’s exact test, Student’s t test, Proportion test, Chi-squared test, Pearson Correlation Coefficient, and Spearman Correlation Coefficient, with a p-value < 0.05 considered statistically significant. Survival comparisons were quantified using log rank Mantel-Cox test and Cox proportional hazards model. Quantitative spatial analysis and cell phenotypic classification for IHC/mIHC were performed using the latest version of QuPath (v0.7.0), where automated cell detection and intensity-based thresholding were applied to calculate the density and distribution of specific cellular subsets.

## Supplementary Table Legends

**Table S1. Summary of Datasets and Patient Clinicopathological Metadata, Related to Figure 1 and Figure S1.** Catalog of the integrated Atlas cohort demographics, detailed by sequencing modalities, sample counts, and treatment stratification across 12 independent single-cell/nucleus RNA-seq datasets (encompassing 16 primary gliomas and 157 secondary brain metastases).

**Table S2. Differentially Expressed Genes (DEGs) for All Cell Subpopulations, Related to Figures 1, 2, 3, 4, and 5.** Comprehensive registry of cluster-specific marker genes, including statistical significance metrics (log2 fold-change, p-value, and adjusted p-value), and baseline expression fractions resolved across all major malignant, myeloid, lymphoid, stromal, and vascular cell lineages.

**Table S3. Summary of Published Gene Signatures for Cell-State Benchmarking and Validation, Related to Figure 4, Figures S2, S3, and S5.** Compendium of reference gene expression signatures extracted from landmark BrM and pan-cancer atlases, compiled to quantitatively evaluate and validate the cell-state annotations of malignant, myeloid, endothelial, and stromal compartments via AUCell scoring.

**Table S4. Patient Clinicopathological Metadata for scRNAseq, IHC/mIHC and CosMx SMI cohorts, Related to Figures 2, 4, 6 and S4.** Comprehensive clinical registry of the validation cohorts, documenting patient demographics, tumor characteristics, and precise tissue allocation tracking for IHC/mIHC and CosMx SMI 6k high-plex profiling.

**Table S5. Downsampled and Balanced Spatial Niche Assignment Matrix from CosMx High-Plex Imaging, Related to Figure 6 and Figure S6.** Cellular neighborhood intensity, coordinate mapping, and local cell-type composition matrix, optimized via computational downsampling and balancing, used to construct and define spatial architectural domains across integrated clinical tissue specimens.

**Table S6. Differential Expression Analysis Across Spatial Niches within Mesenchymal-Like Domains, Related to Figures 6, 7, and S7.** Statistical outputs and differential transcriptomic profiles characterizing the localized microenvironmental features and unique signature genes across the four identified spatial niches (S1, S2, S3, and S4) resolved within the MES-like malignant cell domains.

**Table S7. Proteogenomic Landscapes, Subtype Stratification, and Prognostic Profiles of the Bulk Clinical Cohorts, Related to Figure 7 and Figure S7.**

Comprehensive registry and deconvolution analysis of the bulk transcriptomic and proteomic clinical validation cohorts (n = 312 transcriptomic BrMs; n = 107 independent proteomic BrMs). This table documents the patient clinical metadata, assigned niche-derived molecular subtypes (Proliferative, Angiogenic, Stroma-Reactive, and Quiescent), lineage-specific distribution tracks, and alignment mapping against established consensus BrMS classifications (BrMS1–BrMS4).

## Data and Code Availability

### Data availability

The single-cell and spatially resolved transcriptomic datasets generated in this study have been deposited at Zenodo and are publicly available as of the date of publication at https://zenodo.org/records/20629264. This repository contains: The standard expression files (barcodes, features, and count matrices) for the seven scRNA-seq samples; Sample-specific Seurat v5 objects for the CosMx SMI high-plex spatial transcriptomics data, fully annotated with global cell-type identities; Sample-specific Seurat v5 objects for the CosMx SMI data focused exclusively on the MES-like malignant domains, fully annotated with the four spatial architectural niches (S1–S4). Primary raw sequencing data for the public cohorts integrated into our master single-cell atlas were sourced from open-access repositories (including GEO, EGA, and SRA); specific accession numbers, cohort demographics, and sample metadata are comprehensively tabulated in **Table S1**.

### Code availability

The computational architecture, preprocessing pipelines, and downstream analytical workflows utilized in this study mirror the standardized methodologies established in our previous glioblastoma framework (GRIT-Atlas). All original code, consensus NMF decomposition scripts, and spatial neighborhood analysis pipelines are fully documented and publicly accessible at GitHub: https://github.com/Dr-fly/GRIT-Atlas.

## Acknowledgements

Not applicable.

## Author contributions

**Conceptualization**, Fei Wang, Dengfeng Lu, Zhouqing Chen, Xiaoou Sun, and Zhong Wang; **Methodology**, Fei Wang, Chen Yang, Yanbo Yang, Dengfeng Lu, and Xiaoou Sun; **Software**, Fei Wang, Chen Yang and Run Huang; **Formal analysis**, Fei Wang, Chen Yang, Run Huang, and Yanbo Yang; **Investigation**, Fei Wang, Xiaodong Pang, Chen Yang, Run Huang, Wenqian Cao, Yuhan Bai, Haohao Qiu, Juyi Zhang, Bixi Gao, Chao Ma, and Shengkai Yang; **Data curation**, Fei Wang, Xiaodong Pang, Chen Yang, Run Huang, Wenqian Cao, and Yuhan Bai; **Validation**, Fei Wang, Xiaodong Pang, Wenqian Cao, Yuhan Bai, Haohao Qiu, and Chuanyong Mu; **Resources**, Fei Wang, Xiaodong Pang, Haohao Qiu, Juyi Zhang, Bixi Gao, Chao Ma, Shengkai Yang, Chuanyong Mu, Dengfeng Lu, and Zhong Wang; **Visualization**, Fei Wang and Chen Yang; **Writing – original draft**, Fei Wang, Chen Yang, and Zhouqing Chen; **Writing – review & editing**, Fei Wang, Yanbo Yang, Dengfeng Lu, Zhouqing Chen, Xiaoou Sun, and Zhong Wang; **Supervision**, Dengfeng Lu, Zhouqing Chen, Xiaoou Sun, and Zhong Wang; **Project administration**, Dengfeng Lu, Zhouqing Chen, Xiaoou Sun, and Zhong Wang; **Funding acquisition**, Dengfeng Lu, Zhouqing Chen, Xiaoou Sun, and Zhong Wang.

## Supplementary Figure Legends

**Figure S1.**
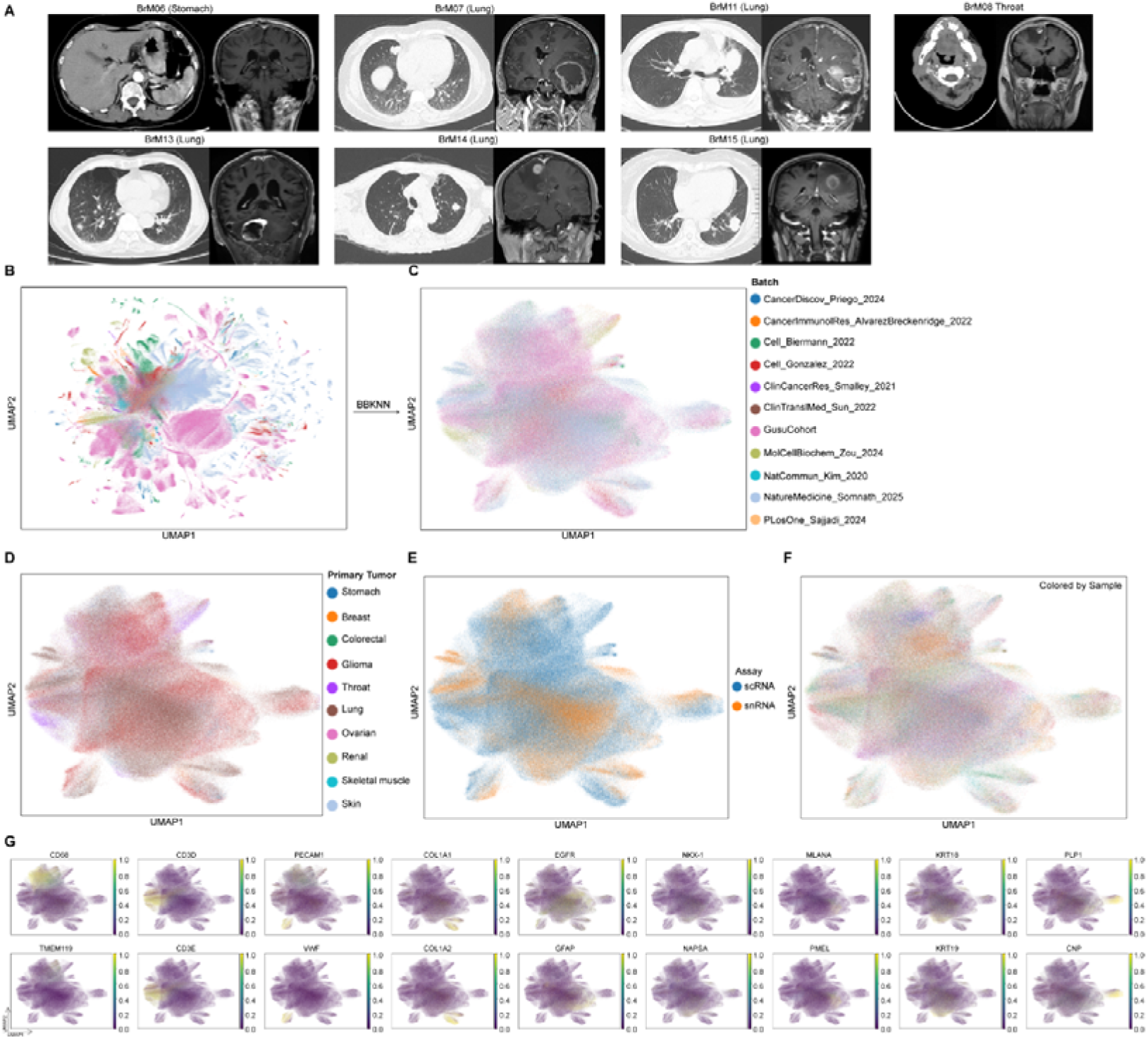
Clinical Imaging, Batch Mitigation, and Multi-dimensional Validation of the Unified Cohort. **(A)** Matched visceral CT images and contrast-enhanced brain MRI scans from seven patients within the in-house cohort. **(B–C)** Global single-cell UMAP embeddings before **(B)** and after **(C)** batch effect correction using the BBKNN algorithm. **(D–F)** Integrated UMAP embeddings colored by primary tumor group **(D)**, center of origin **(E)**, and sequencing modality **(F)** to confirm technical variation mitigation. **(G)** Two-row feature plot matrix displaying single-cell transcript expression of canonical marker genes across the six major cell lineages.

**Figure S2.**
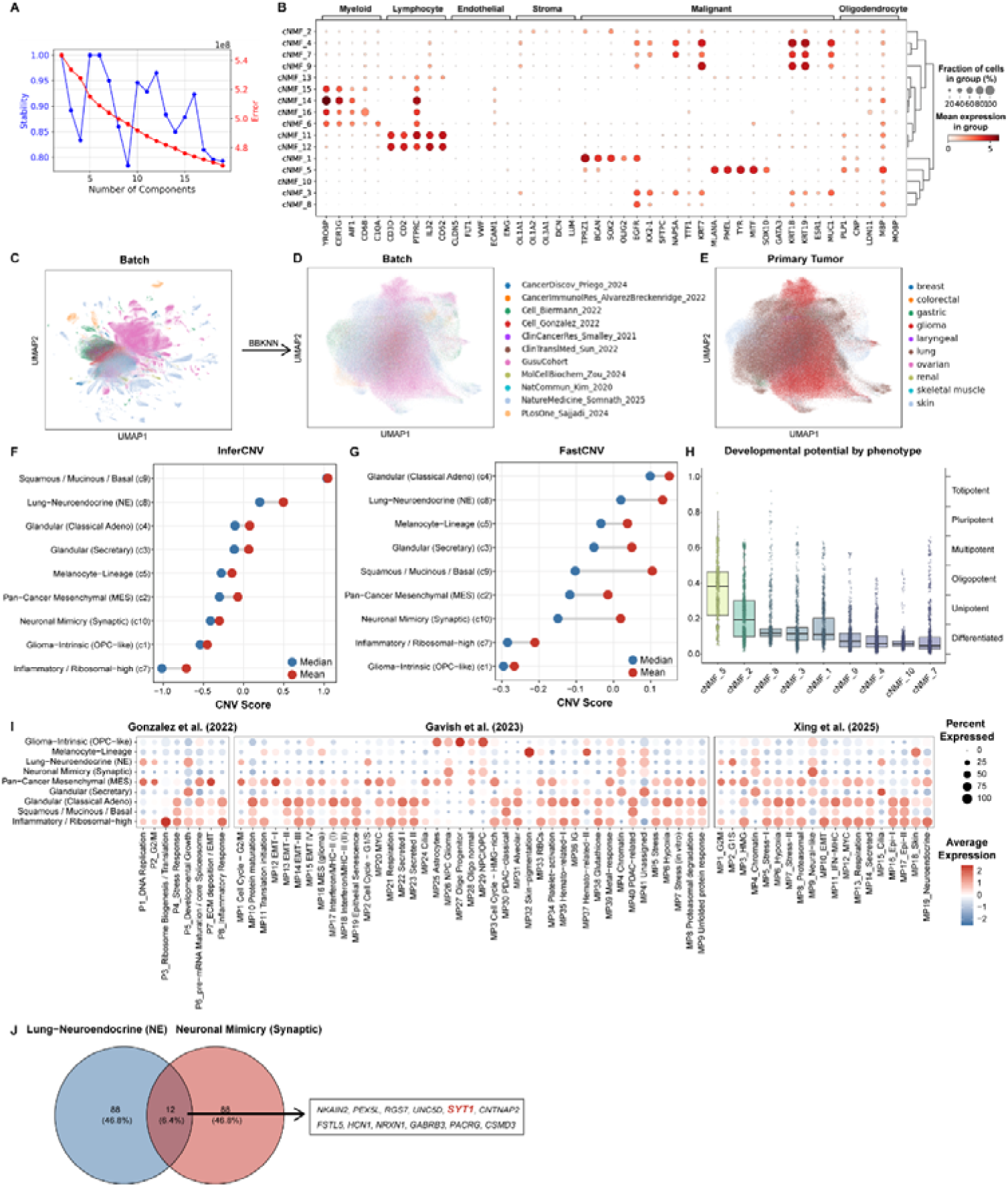
Quality Control, Batch Correction, and Multi-dimensional Validation of Malignant cNMF Programs. **(A)** Evaluation of cNMF model stability and reconstruction errors across varying factorization ranks (K). **(B)** Marker gene dot plots charactering four excluded noise or confounding cNMF components. **(C–E)** UMAP embeddings colored by batch **(C, D)** and disease group **(E)** confirming integration consistency. **(F–G)** Chromosomal instability (CIN) profiling across cNMF programs via inferCNV **(F)** and fastCNV quantification scores **(G)** (blue dot, median; red dot, mean). **(H)** CytoTRACE2 differentiation potential scores mapped onto UMAP embedding to track stemness. **(J)** Intersection analysis showing 12 overlapping marker genes between sub-clusters c8 and c10.

**Figure S3.**
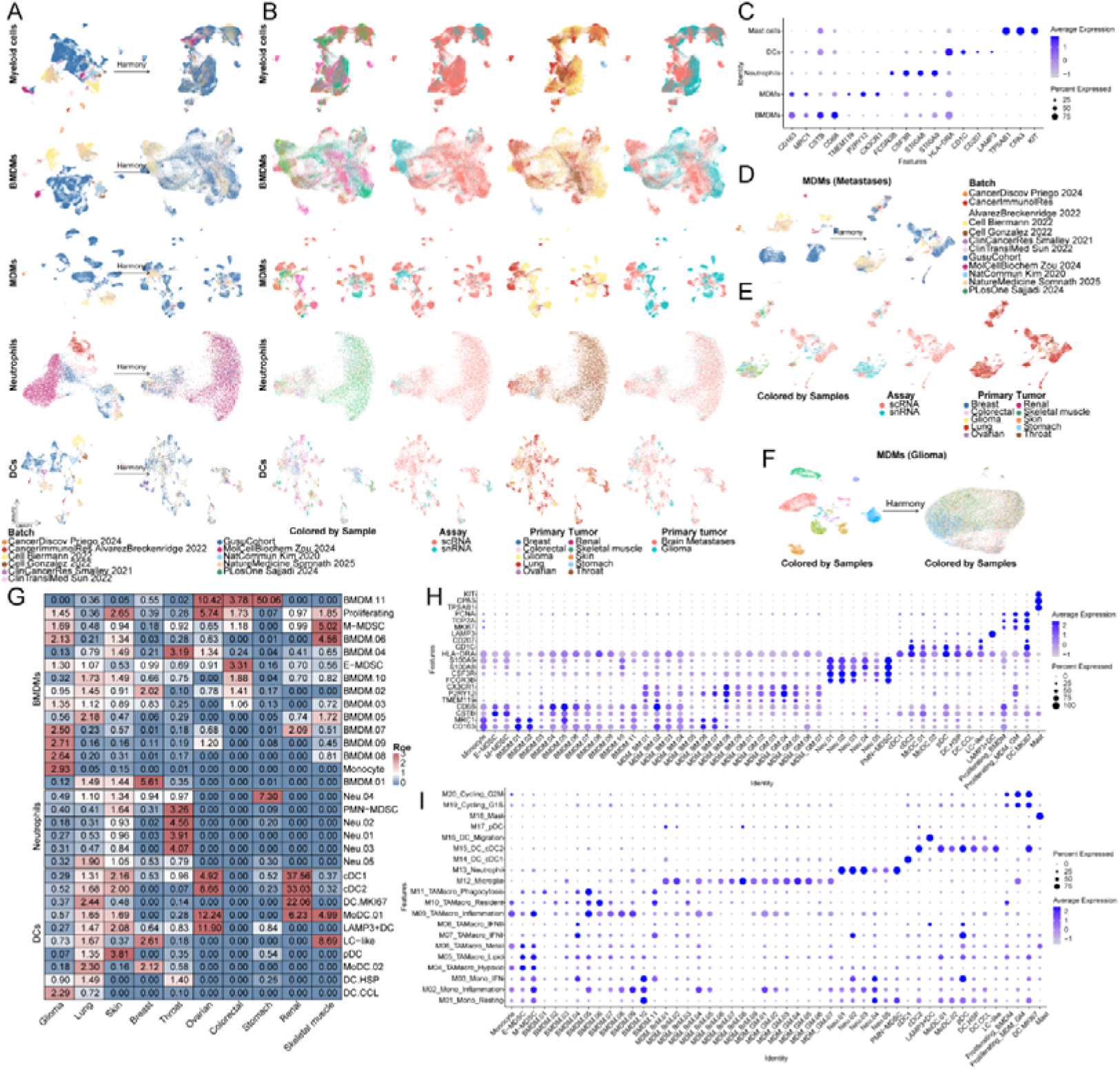
Multi-dimensional Quality Control, Batch Mitigation, Sub-clustering Optimization, and Reference Benchmarking of the Myeloid Landscape. **(A)** Myeloid UMAP embeddings before (left) and after (right) Harmony integration. **(B)** Myeloid UMAP panels colored by center of origin, sequencing modality, and primary tumor category confirming batch mitigation. **(C)** Canonical marker expression dot plot defining major myeloid categories. **(D–E)** Target-specific Harmony batch correction **(D)** and integration assessment panels **(E)** across BrM-associated MDMs. **(F)** Glioma-associated MDM UMAP embeddings before (left) and after (right) Harmony correction. **(G)** Observed-to-expected cell ratio (Ro/e) heatmap tracking myeloid sub-cluster tissue preferences across cohorts. **(H)** Dot plot visualizing transcript expression of canonical marker genes across finely resolved myeloid sub-clusters. **(I)** Cross-dataset benchmarking correlating resolved myeloid sub-clusters against established reference archetypes from Xing et al. via AUCell.

**Figure S4.**
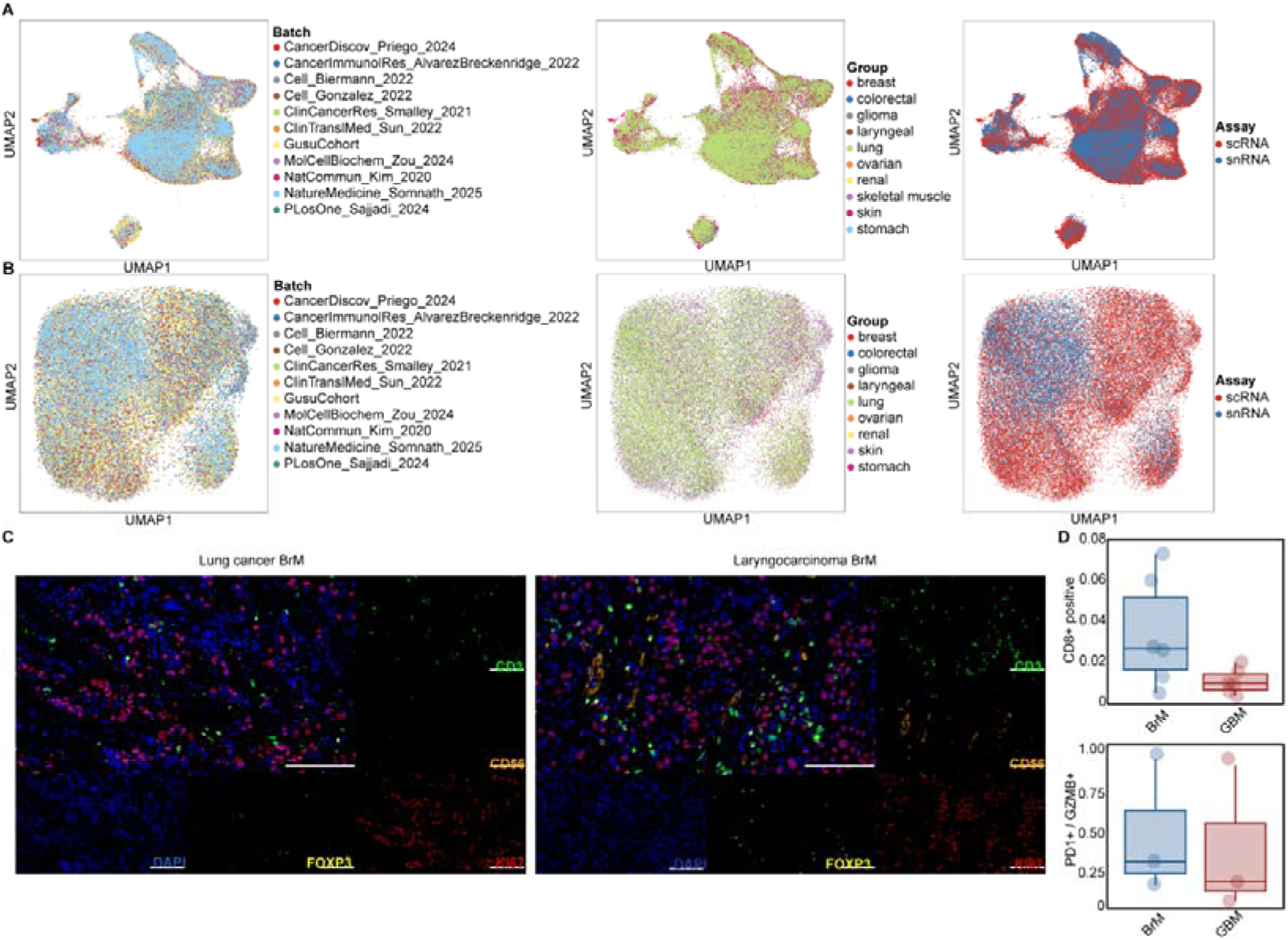
Multi-dimensional Quality Control, Integration Assessment, and Histological Validation of Lymphoid Compartment. **(A)** Global lymphoid UMAP embeddings colored sequentially by batch, disease group, and sequencing assay type. **(B)** T cell sub-clustering UMAP panels colored by batch, disease group, and sequencing assay type. **(C)** Representative mIHC staining profiles (DAPI, blue; CD3, green; CD56, orange; FOXP3, yellow; Ki-67, red) capturing immune phenotypes in lung and laryngocarcinoma BrM. Scale bar, 130 μm. **(D)** Histological quantification of T cell exhaustion and cytotoxicity across cohorts. Upper box plot shows IHC-based CD8+ cell positivity rates across lung BrM and GBM sections; lower box plot tracks the relative fractions of exhausted (CD3+PD-1+) and cytotoxic (CD3+GZMB+) T cells around cohort medians to validate single-cell functional dichotomy.

**Figure S5.**
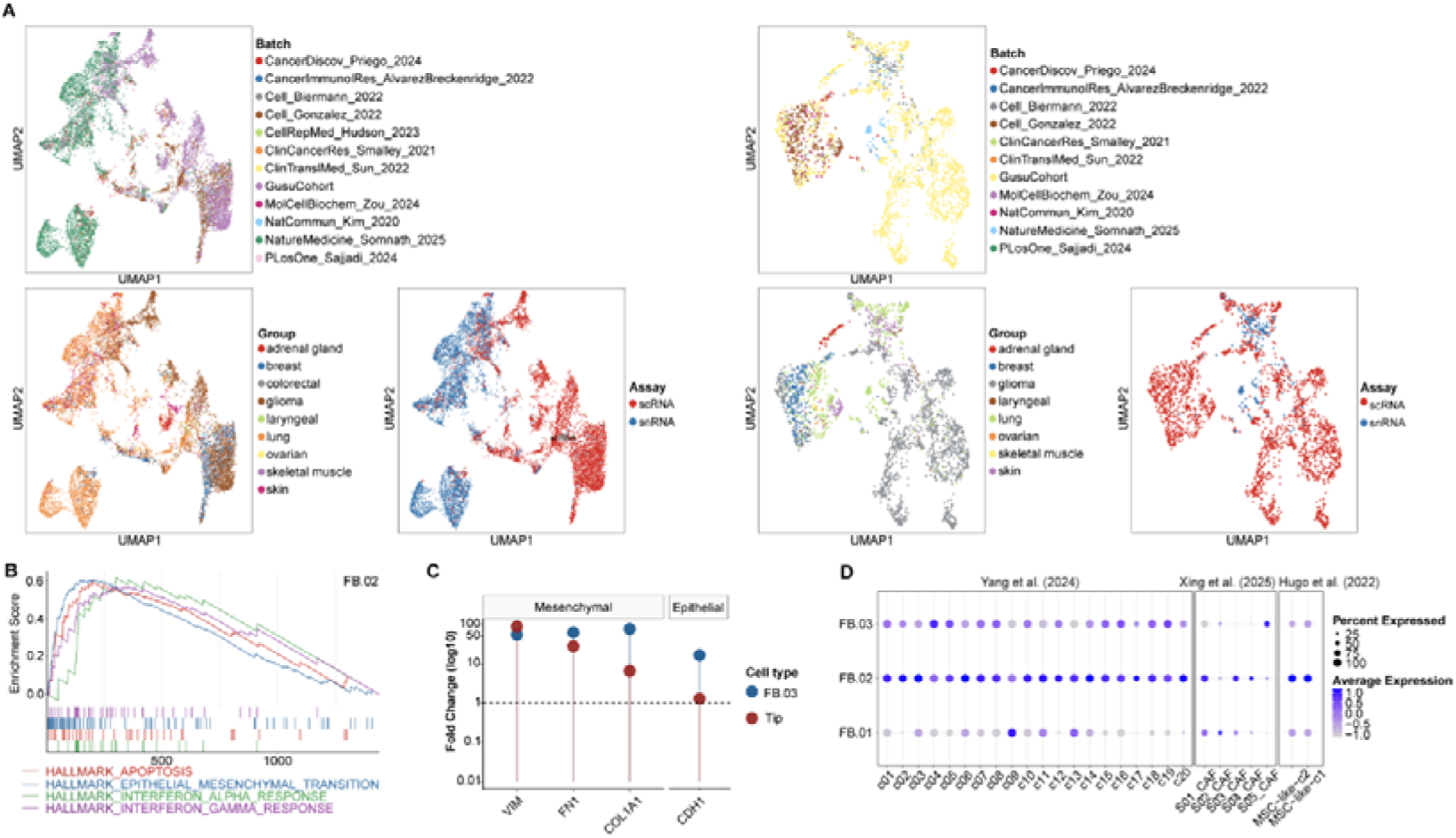
Integration Assessment and Supplementary Validation of Stromal and Vascular Remodeling Programs. **(A)** Stromal and vascular UMAP embeddings colored by batch, disease group, and sequencing assay type to confirm integration stability. **(B)** Supplementary GSEA curves showing representative functional pathway programs associated with FB.02.COL1A1. **(C)** Lollipop plot comparing EMT-related marker expression between FB.03.MMP14 and Tip.PLVAP to contrast matrix-degrading and pro-invasive states. **(D)** Cross-dataset signature benchmarking comparing FB.01.PDGFRA, FB.02.COL1A1, and FB.03.MMP14 against independent brain metastasis stromal datasets to confirm external consistency.

**Figure S6.**
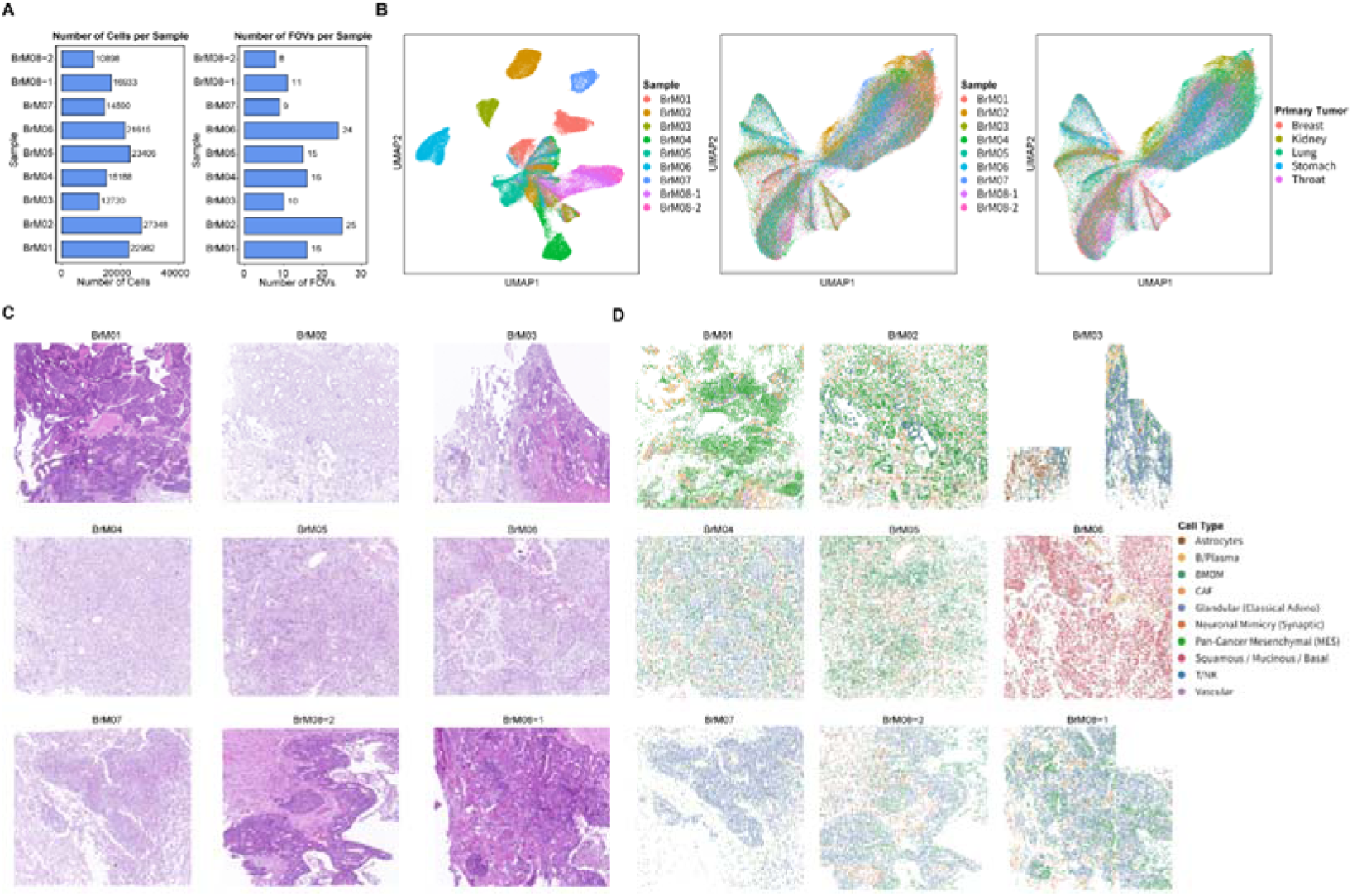
Quality Control Metrics, Integration Assessment, and Histological Architecture of the CosMx Spatial Cohort. (A–B) Bar plots showing the total number of single cells recovered after quality control filtering **(A)** and fields-of-view (FOVs) analyzed **(B)** per individual sample. **(C–E)** Global spatial UMAP visualizations before **(C)** and after **(D, E)** batch correction, colored by sample **(C, D)** and primary tumor site of origin **(E)**. **(F)** Representative serial H&E sections capturing global tissue architecture across the 9 specimens from 8 patients. **(G)** Spatial distribution maps of annotated cell lineages corresponding exactly to the tissue coordinates shown in **(F)**.

**Figure S7.**
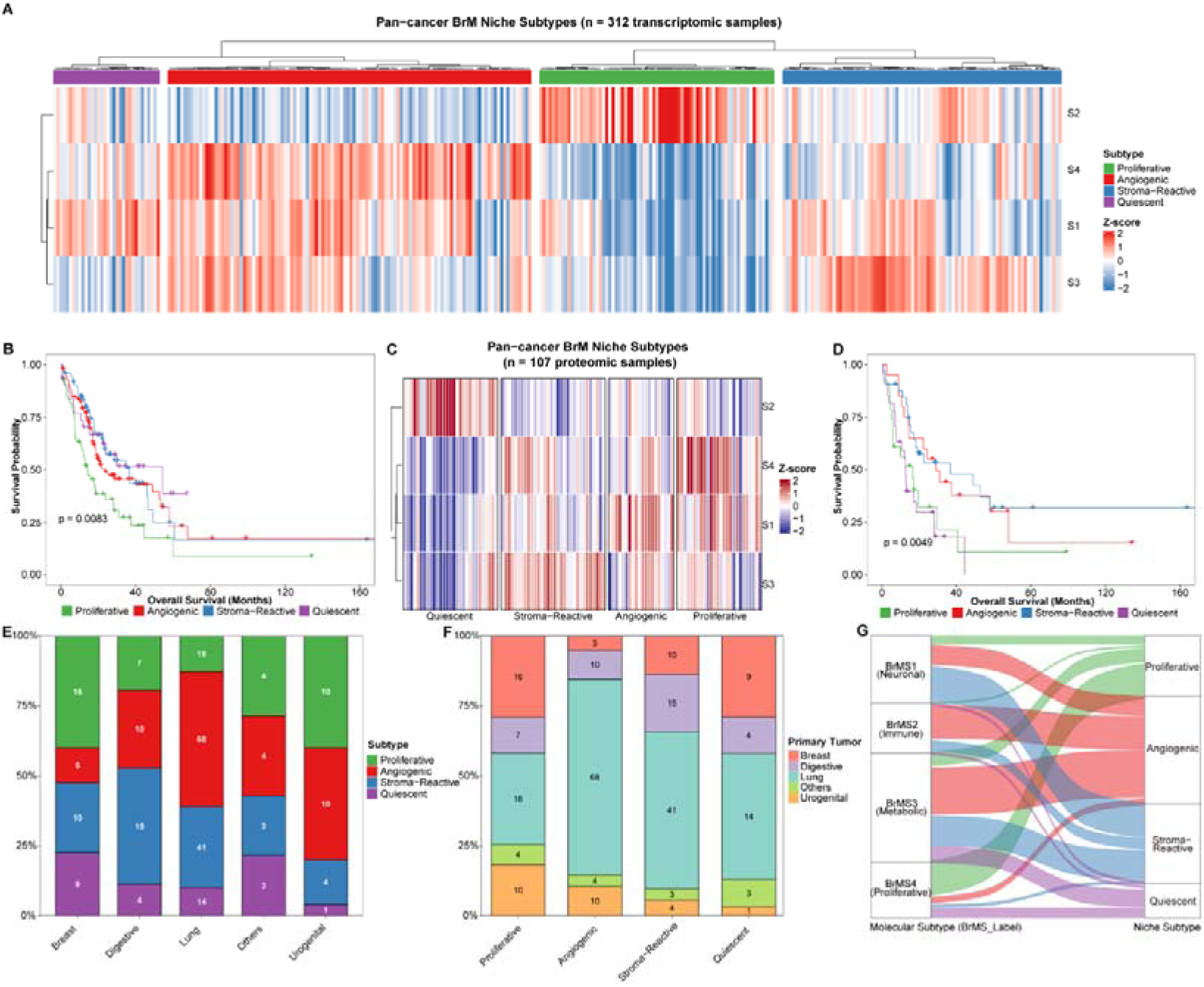
Proteogenomic Landscape, Lineage Preferences, and Prognostic Stratification of Niche-Defined Clinical Subtypes. **(A)** Unsupervised hierarchical clustering heatmap of row-standardized niche activity scores (S1–S4) within the bulk transcriptomic cohort (n = 314), partitioning patients into four clinical subtypes. **(B)** Kaplan-Meier overall survival curves of BrM patients stratified by the four transcriptomic niche-defined clinical subtypes (n = 314). P-value calculated by log-rank test. **(C)** Unsupervised hierarchical clustering validation heatmap of row-standardized protein activity scores within the validation proteomic dataset (n = 107). **(D)** Kaplan-Meier overall survival curves of BrM patients stratified by the four proteomic niche-defined clinical subtypes (n = 107). P-value calculated by log-rank test. **(E–F)** Stacked bar plots illustrating percentage distributions of niche-defined subtypes across primary tumor groups **(E)** and primary tumor groups across clinical subtypes **(F)**, with absolute sample counts indicated inside segments. **(G)** Alluvial diagram mapping sample overlap and classification alignment between established molecular BrMS subtypes (BrMS1 to BrMS4) and the four niche-defined clinical subtypes.

